# Hypercholesterolemia risk associated GPR146 is an orphan G-protein coupled receptor that regulates blood cholesterol level in human and mouse

**DOI:** 10.1101/2020.01.09.901041

**Authors:** Fangfang Han, Xiao Liu, Chuanfang Chen, Yinan Liu, Mingkun Du, Yangyang Guan, Yiliang Zhang, Dehe Wang, Musaddeque Ahmed, Xuedan Li, Xiaomin Liu, Yuxian Wu, Yu Zhou, Yong Liu, Bao-Liang Song, Housheng Hansen He, Yan Wang

**Author notes:** These authors contributed equally (Fangfang Han, Xiao Liu, Chuanfang Chen, Yinan Liu). Corresponding author: Yan Wang.

## Abstract

Genome-wide association studies (GWAS) have identified hundreds of genetic variants associated with dyslipidemia. However, about 95% of of these variants are located in genome noncoding regions and cluster in different loci. The disease-causing variant for each locus and underline mechanism remain largely unknown. We systematically analyzed these noncoding variants and found that rs1997243 is the disease-causing variant in locus 7p22, which is strongly associated with hypercholesterolemia. The rs1997243 risk allele is associated with increased expression of *GPR146* in human and targeted activation of the rs1997243 site specifically up regulates *GPR146* expression in cultured cells. GPR146 is an orphan G-protein coupled receptor that is located on plasma membrane and responses to stimulation of heat-inactivated serum. Disrupting *gpr146* specifically in the liver decreases the blood cholesterol level and prevents high-fat or high-fat high-cholesterol diets induced hypercholesterolemia in mice. Thus we uncovered a novel G-protein coupled receptor that regulates blood cholesterol level in both human and mouse. Our results also suggest that antagonizing GPR146 function will be an effective strategy to treat hypercholesterolemia.

---

Genome-wide association study (GWAS) is a powerful tool to ascertain the contribution of common genetic variants in population-wide diseases variability. Hundreds of GWAS studies have been applied to a variety of diseases or traits including dyslipidemia, diabetes, and hypertension et al^1^. More than 93% of these disease- and trait-associated variants are located in the non-coding regions, makes it difficult to evaluate their function^2^. Previous studies showed that these disease- and trait-associated variants are concentrated in regulatory DNA, with about 80% of all noncoding GWAS single nucleotide polymorphisms (SNPs) or linkage disequilibrium (LD) SNPs are located within DNase I hypersensitive sites (DHS)^2^, which suggest that most of these noncoding SNPs function through transcriptional regulation.

Hypercholesterolemia is the leading risk factor for cardiovascular diseases. Current evidence suggests that the heritability for blood cholesterol level is high, with 40-50% for low-density lipoprotein cholesterol (LDL-C) and 40-60% for high-density lipoprotein cholesterol (HDL-C)^3,4^. GWAS has been performed extensively on blood lipids traits and more than 300 risk loci were found in different populations^5^. These loci cover almost all the well-known genes that are important in lipid metabolism, such as *LDLR^6^*, *ARH^7^*, *ABCA1^8^*, *ABCG5/G8^9^*, *PCSK9^10^*, *NPC1L1^11,12^*, *LIMA1^13^*, et al. Although GWAS is a powerful approach, we find that about 2/3 of these risk loci are located in noncoding regions and are not close to any gene known plays a role in lipid metabolism. The disease-causing variants in these loci and their underlying molecular mechanism remain largely unknown, which prevent the interpretation of the GWAS results and their application in precise medicine. On the other hand, these noncoding regions may harbor novel genes or signaling pathways involved in lipid metabolism and provides a valuable resource for further mechanistic studies.

We systematically analyzed these noncoding loci and primarily focused on loci that are not close to any gene known plays a role in lipid metabolism. One such locus 7p22 is strongly associated with increased level of blood cholesterol in multi cohorts^14–16^ (Fig S1a, S2b). The lead SNP rs1997243 is a common variant and has the highest frequency in European population but is absent in East Asian (Fig S1b). It is located in the noncoding region and has a strong linkage disequilibrium non-synonymous variant rs11761941 (*GPR146* p. Gly11Glu) in some populations (Fig S1a, S2a, S2b). Both rs1997243 and rs11761941 are significantly associated with blood cholesterol level (Fig S2b)^15^. However, the *GPR146* p. Gly11 is not conserved and has been substituted with Asp, Asn or Ala in many other species except *Gray wolf* (Fig S2c). Bioinformatics analysis also predicts that *GPR146* p. Gly11Glu is benign and neutral (Fig S2d), rendering rs11761941 less likely to be the disease-causing variant.

We reasoned that any SNPs that have strong linkage disequilibrium with the lead SNP could be the real disease-causing variant. Since all these variants are located in the noncoding region, the real disease-causing variant is most likely located in the regulatory region, such as regions marked by DNase I hypersensitivity and/or histone modifications marker H3K27ac and H3K4me3^17–21^. We systematically analyzed all SNPs that have strong linkage disequilibrium with the lead SNP rs1997243 in 7p22 locus. There are 125 SNPs were identified, with 28 of them are located in genome active regions (Fig S3a, S3i, Table S1). We then applied luciferase reporter assay to compare the transcriptional activities between the minor allele and the major allele for each of these SNPs. Promoter sequence of *APOA1* was used as a positive control for the assay (Fig S3b). We found that across all SNPs tested, only rs1997243 shows increased promoter activity compared with its reference allele under similar transfection efficiency (Fig 1b, Fig S3c-k). The rs1997243 does not change enhancer activity in enhancer luciferase reporter assay (Fig 1c, Fig S3l), which is consistent with enriched promoter specific histone marker H3K4me3 at this position (Fig 1a)^17,20,21^.

**Figure 1.**
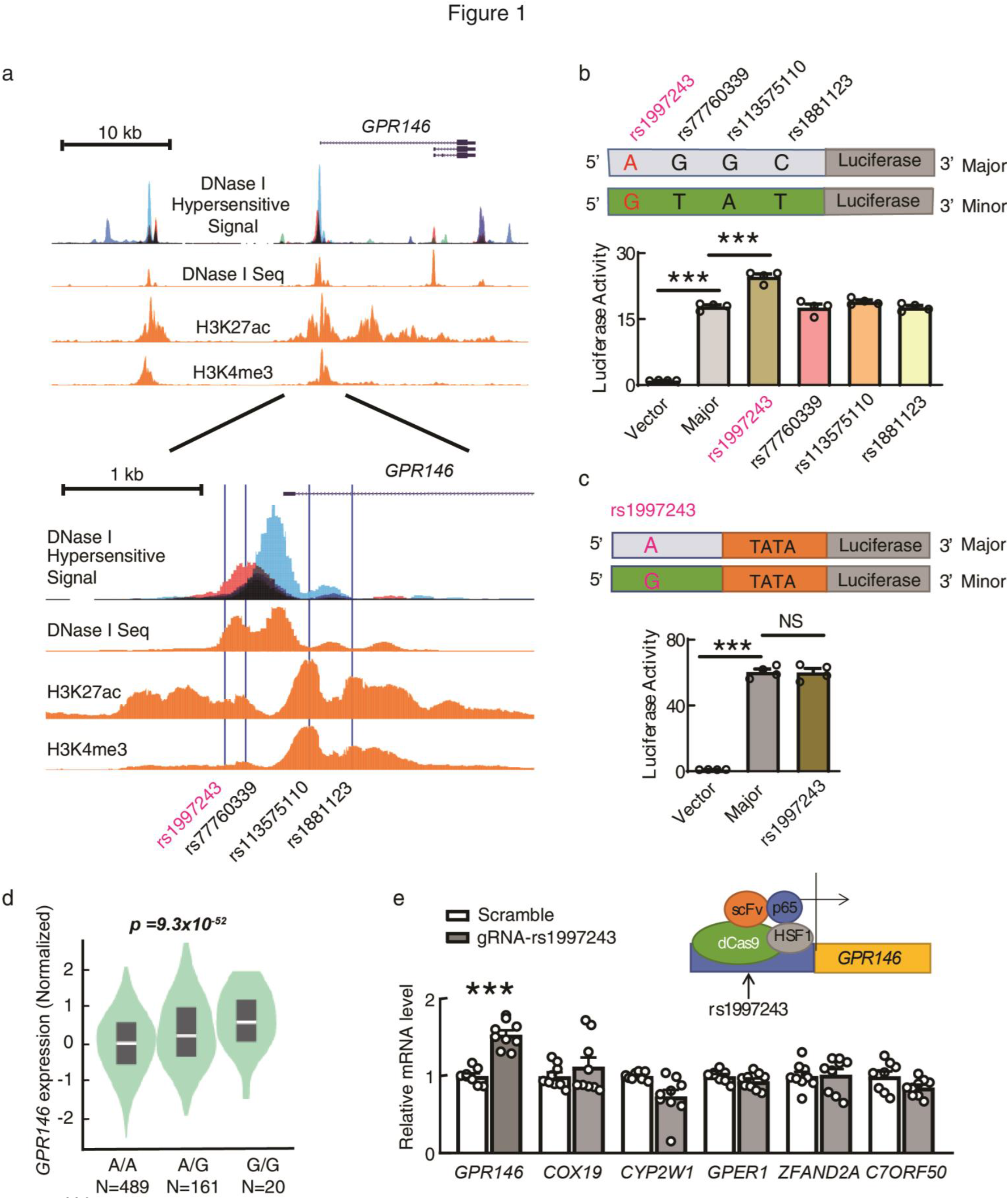
rs1997243 is the disease-causing variant in 7p22 locus. **a**, Genome active regions in 7p22 locus. Genome active regions are markered by DNase I hypersensitive signal from 95 cell lines and DNase I seq, H3K27ac Chip-seq, H3K4me3 Chip-seq signals from human liver samples. Data were downloaded from ENCODE and plotted with UCSC Genome Brower as described in Methods. The lead SNP rs1997243 and three other SNPs that have strong linkage disequilibrium with rs1997243 are located in one of the genome active region close to the transcriptional starting site of *GPR146*. **b**, rs1997243 minor allele increases the promoter activity of the genome sequence. Genome sequence covering different SNPs was cloned into upstream of luciferase reporter gene. Plasmid with reference allele (Major) or each of the minor allele (Minor) was transfected into HepG2 cells separately. The luciferase activity was assayed as described in Methods. Luciferase activity in the vector-transfected cells was set to 1. **c**, rs1997243 does not change the enhancer activity of the genome sequence. Genome sequence covering the reference allele (Major) or rs1997243 risk allele (Minor) was cloned into upstream of TATA box mini-promoter followed by luciferase reporter gene. The luciferase activity was assayed and analyzed exactly the same as in **b**. **d**, rs1997243 G-allele is significantly associated with increased expression of *GPR146* in human. eQTL analysis for rs1997243 was performed in 670 human whole blood samples as described in Methods. **e**, Transcriptional activation of rs1997243 site increases *GPR146* expression in HepG2 cells. The transcription activator complex (scFV, p65, HSF1) was targeted to rs1997243 site through enzymatic dead Cas9 (dCas9) together with a specific gRNA sequence to rs1997243 position in HepG2 cells. Gene expression levels were assayed by RT-PCR analysis. All data are expressed as means ± SEM and *p* values were calculated using Student’s test (\*\*\**p*<0.001).

Expression quantitative trait loci (eQTL) analysis showed that the rs1997243 minor allele (G-allele) is strongly associated with increased expression of *GPR146* in human (Fig 1d). Targeted activation of rs1997243 site with enzymatic dead Cas9 (dCas9) system and a gRNA specifically to the rs1997243 position increases the expression level of *GPR146* significantly, with no detectable impact on other genes in this region (Fig 1e, Fig S2e). These data suggest that the rs1997243 is the disease-causing variant and may increase blood cholesterol level through up regulating *GPR146* expression.

GPR146 is an orphan G-protein coupled receptor that is highly expressed in liver and adipose tissue of both human and mouse (Fig S4a, Fig 2a). In the liver, it specifically expresses in hepatocytes (Fig 2b). By prediction, it contains typical seven transmembrane domains with N terminal facing extracellular compartment (Fig S4b, c). When expressed in cells, GPR146 is located in membrane fraction and is located on plasma membrane, which suggests that it may function as a receptor (Fig 2c, d). GPCRs typically signaling through Gα_s_, Gα_i/o_, Gα_q/11_, Gα_12/13_ or Gβ/γ and regulate cAMP production, intracellular Ca^2+^ mobilizations, ERK/MAPK activity or small G protein RhoA activity et al^22^. We found that GPR146 responses to serum filtered by 3 kDa cut-off filter and activates the transcriptional activity of cAMP response element (CRE) (Fig 2e). Moreover, this response is preserved when the serum was further heat-inactivated by boiling and can be fully blocked by PKA inhibitor H-89 (Fig 2f). Taken together, our data suggests that GPR146 is a cell signaling receptor that responses to serum stimulation and activates the PKA signaling pathway.

**Figure 2.**
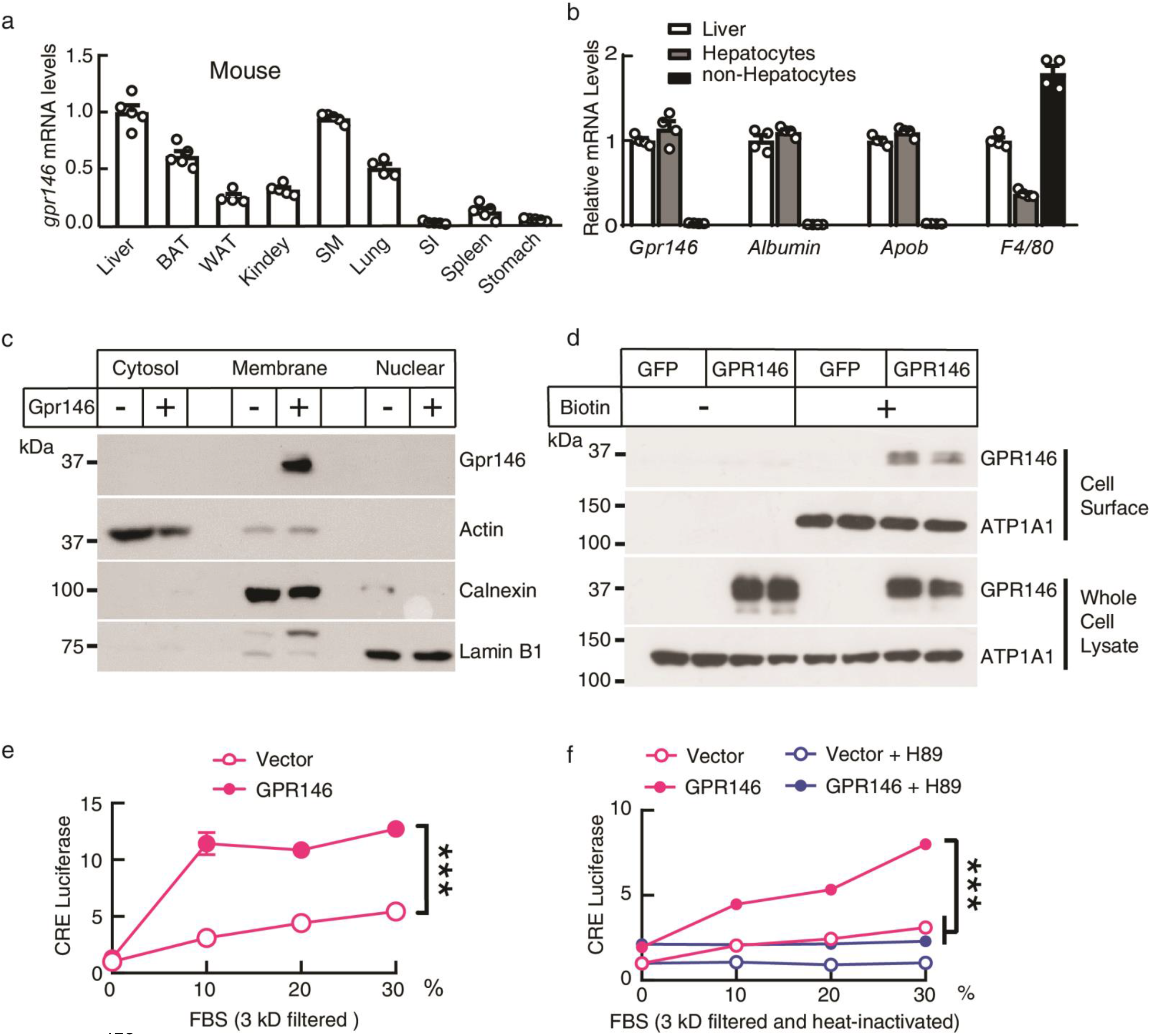
GPR146 localizes on plasma membrane and responses to serum stimulation. **a**, Tissue distribution of *gpr146* in mice. C57BL/6J wild type mice were fast overnight and indicated tissues were collected for RT-PCR analysis. Expression levels were normalized to liver which was set to 1 (N=5, male, 8 weeks). BAT, brown adipose tissue; WAT, white adipose tissue; SM, skeletal muscle; SI, small intestine. **b**. *Gpr146* is specifically expressed in hepatocytes of mouse liver. Primary hepatocytes and non-hepatocytes are separated and subjected to RT-PCR analysis as described in Methods. *Albumin* and *Apob* are hepatocytes marker genes, *F4/80* is a macrophage marker gene. **c**. Gpr146 is located in membrane fraction. Flag tagged Gpr146 was expressed in 293T cells by transient transfection. Cytosol, membrane and nuclear fractions were isolated as described in Methods. Gpr146 was detected with anti-Flag antibody. Actin, Calnexin and Lamin B1 were used as markers for each fraction. **d**, GPR146 is located on plasma membrane. GPR146 tagged with Flag was expressed in 293T cells by transient transfection. Cell surface fraction was isolated with biotinylation in part of the cells as described in Methods. GPR146 was detected with an anti-GPR146 polyclonal antibody. ATP1A1 was used a cell surface marker. **e**, GPR146 responses to 3 kD filtered serum and activates cAMP response element (CRE) activity. HEK293 cells expressing empty vector or GPR146 together with CRE-Luciferase reporter were treated with indicated amount of fetal bovine serum (FBS) that has passed through the 3 kD cut-off filter. Luciferase activities were measured accordingly. **f**, GPR146 responses to heat-inactivated serum and actives CRE activity through PKA pathway. HepG2 cells expressing empty vector or GPR146 together with CRE-Luciferase reporter were treated with indicated amount of FBS that has passed through the 3 kD cut-off filter and further heat-inactivated by boiling. Part of the cells was treated in the presence of PKA inhibitor H-89 (10 μm). All data are expressed as means ± SEM and *p* values were calculated using Student’s test (\*\*\**p*<0.001). All experiments were repeated at least twice with similar results.

To further study its function *in vivo*, we generated *gpr146* knockout mouse model with *Cre*-*LoxP* system (Fig S5a). Totally we got 6 lines of *LoxP* positive F1 mice and they were genotyped by genome sequencing (data not shown), southern blot analysis (Fig S5b) and PCR genotyping (Fig S5c). Line 92 was used for all experiments except otherwise indicated. We generated whole body, liver specific and adipose tissue specific *gpr146* knockout mice by crossing the *LoxP/LoxP* mice with Cre recombinase driven by *CMV*, *albumin* and *adiponectin* promoters respectively. Whole body *gpr146*^−/−^ mice have significantly decreased blood cholesterol level compared with their littermate controls (Fig S6a, b). In contrast, the adipose tissue specific *gpr146*^−/−^ mice have no detectable difference of blood lipids levels compared with their littermate controls (Fig S6c, d). The liver specific *gpr146*^−/−^ (Li-*gpr146*^−/−^) mice have significantly decreased blood cholesterol level as the whole body *gpr146*^−/−^ mice and are protected from high-fat high-cholesterol diet induced hypercholesterolemia (Fig 3a-d), which suggest that gpr146 regulates blood cholesterol level mainly through the liver. Consistent with decreased plasma cholesterol level, both ApoB-100 and ApoB-48 protein levels are significantly decreased in the plasma, especially under high-fat high-cholesterol diet feeding (Fig 3e, f). ApoA1 is also slightly decreased in Li-*gpr146*^−/−^ mice under high-fat high-cholesterol diet feeding (Fig 3e, f). Moreover, Li-*gpr146*^−/−^ mice are protected from high-fat diet induced hypercholesterolemia (Fig 4a). To test whether acutely suppressing *gpr146* will decrease blood cholesterol level, we knocked down *gpr146* in livers of adult mice through adeno-associated virus mediated shRNA delivering. As shown in Figure 4b, knocking down *gpr146* in the liver significantly decreases the blood cholesterol level, which indicates that blocking *gpr146* function will be an effective strategy to decrease blood cholesterol in adults. These results were confirmed in Li-*gpr146*^−/−^ mice derived from an independent F1 line (Fig 4c, d, Fig S5b). Taken together, our results clearly demonstrate that GPR146 positively regulates blood cholesterol level, which is consistent with increased blood cholesterol level in humans with rs1997243 minor allele.

**Figure 3.**
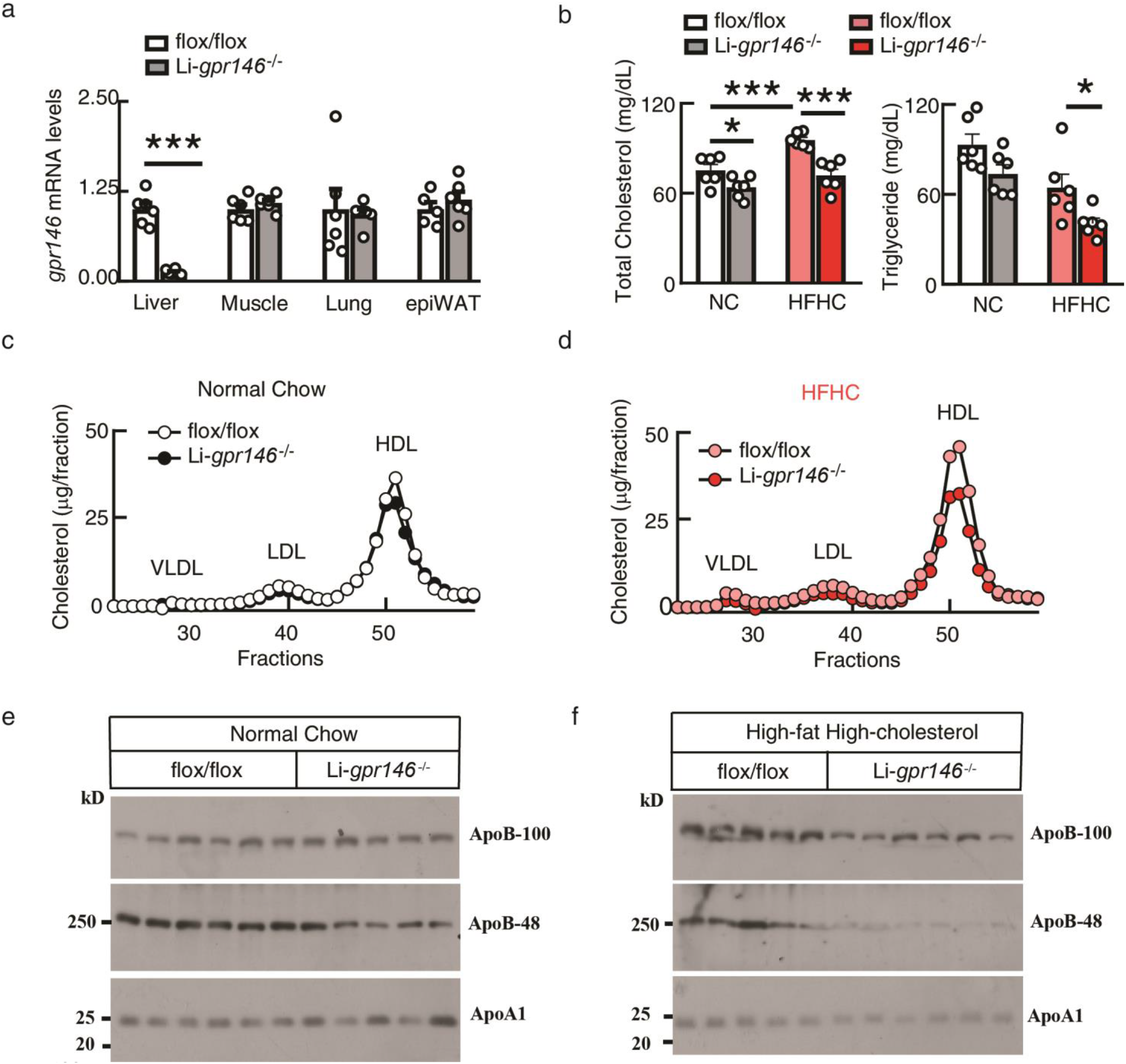
Liver specific *gpr146* knockout mice have decreased blood cholesterol level and are protected from high-fat high-cholesterol diet induced hypercholesterolemia. **a**, *Gpr146* expression levels in liver specific *gpr146* knockout mice (Li-*gpr146*^−/−^) and their littermate controls. Indicated tissues were collected from mice fasted for 3 hours in the early morning and subjected to RT-PCR analysis. Expression levels were normalized to the control group which was set to 1 (N=6, female, 7-8 weeks). epiWAT: epididymal white adipose tissue. **b**, Li-*gpr146*^−/−^ mice have decreased blood cholesterol level compared with their littermate controls. Blood was collected from overnight fasted mice fed with normal chow (NC) or high-fat high-cholesterol (HFHC) for 2 weeks. Plasma levels of total cholesterol and triglyceride were measured as described in Methods (N=6, female, 10-13 weeks). **c**, **d**, Pooled plasma from b was fractioned by Fast Protein Liquid Chromatography. Cholesterol concentration in each fraction was measured accordingly. **e, f**, Plasma levels of ApoB and ApoA1 in Li-*gpr146*^−/−^ and control mice. Plasma from b was separated on SDS-PAGE and subjected to immunoblot analysis as described in Methods. All data are expressed as means ± SEM and *p* values were calculated using Student’s test (\**p*<0.05, \*\*\**p*<0.001). All experiments were repeated at least twice with similar results.

**Figure 4.**
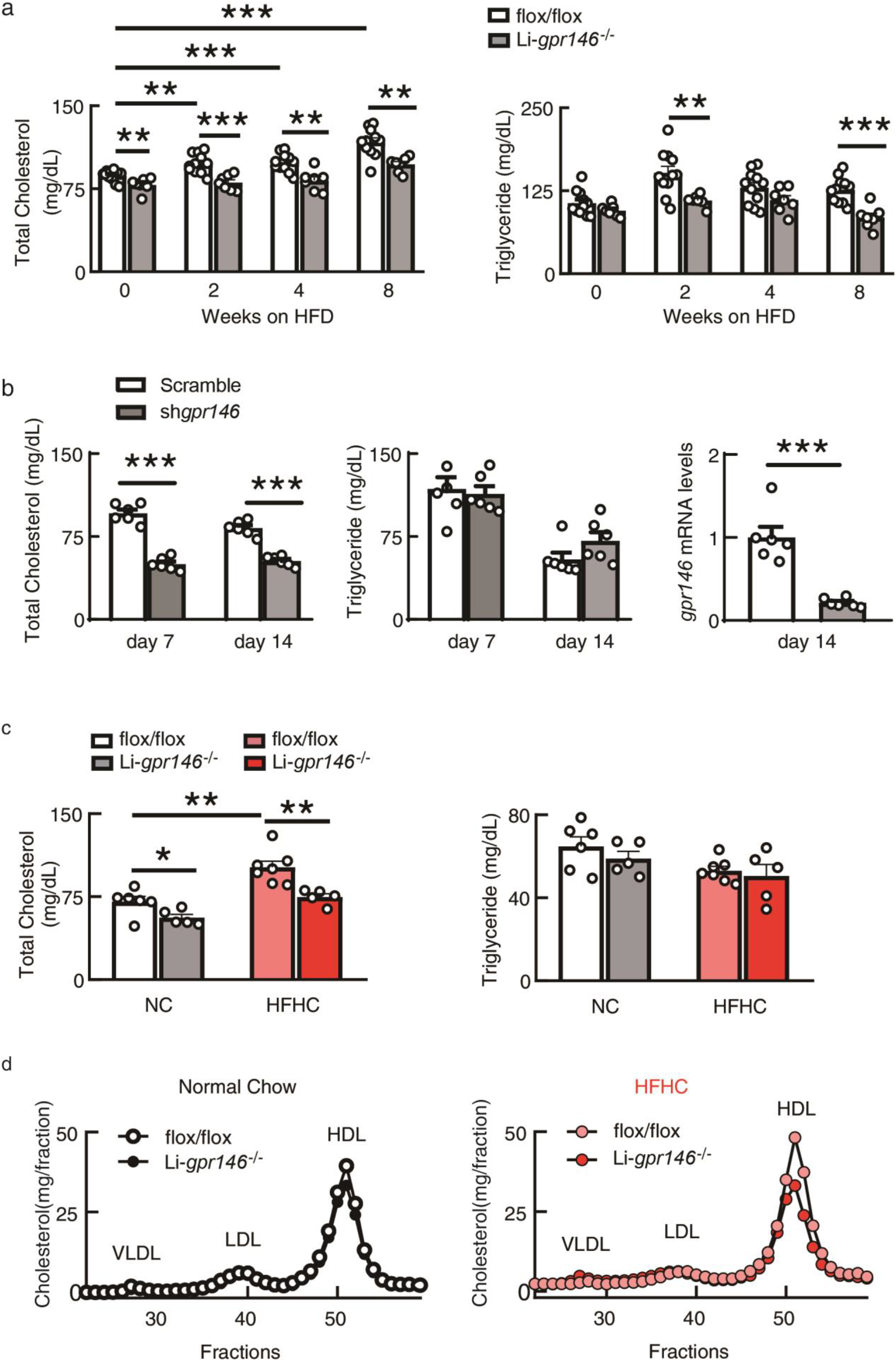
Decreased blood cholesterol level in Li-*gpr146*^−/−^ and *gpr146* knocking down mice. **a**, Li-*gpr146*^−/−^ mice are protected from high-fat diet induced hypercholesterolemia. Mice were fed with high-fat diet (HFD) for eight weeks and blood was collected after overnight fasting at indicated time points. Plasma levels of total cholesterol and triglyceride were measured accordingly (N=7-11, male, 10-12 weeks old when start feeding). **b**, Knocking down *gpr146* in livers of adult mice decreases blood cholesterol level. C57BL/6J wild type mice were infused with adeno-associated virus expressing a scramble or shRNA against *gpr146* through the tail veins. Blood was collected after overnight fasting at one and two weeks after injection. Plasma levels of total cholesterol and triglyceride were measured accordingly. Knocking down efficiency was measured by RT-PCR analysis in liver (right) (N=5, male, 8 weeks). **c**, Plasma levels of cholesterol and triglyceride in Li-*gpr146*^−/−^ mice derived from a second F1 line. The flox/flox mice used in this experiment were derived by crossing line 96 and 97 of F1 mice (Fig S5b). Li-*gpr146*^−/−^ and their littermate controls were fed with normal chow (NC) or high-fat high-cholesterol (HFHC) diet for 2 weeks (n=5-7/group, female, 9-11 weeks). **d**, Pooled plasma from c was fractioned by Fast Protein Liquid Chromatography. Cholesterol concentration for each fraction was measured enzymatically as described in Methods. All data are expressed as means ± SEM and *p* values were calculated using Student’s test (\**p*<0.05, \*\**p*<0.01, \*\*\**p*<0.001). All experiments were repeated at least twice with similar results.

In summary, through bioinformatics analysis and experimental verification we found a noncoding disease-causing variant rs1997243 in locus 7p22. The risk allele of rs1997243 up regulates an orphan G-protein coupled receptor *GPR146* that plays an important role in regulating blood cholesterol level. We believe that the increased expression level of *GPR146* can at least partially explain the disease-causing effect of rs1997243 in human.

In contrast to the causal variants in Mendelian disease, which typically confer large effect, the common variants from GWAS usually have modest effects for each of them. This is especially true for GWAS SNPs that are located in genome noncoding regions. However, variants that explain a small proportion of the traits may provide substantial biological or therapeutic insights. The rs1997243 confers a modest effect on total blood cholesterol level with effect size of 0.033^14^. However, combining bioinformatics analysis and functional studies we found that the downstream target gene *GPR146* has a large impact on blood cholesterol level. Our data also reveal that GPR146 responses to an endogenous ligand in the serum and actives the PKA signaling pathway, which suggest that GPR146 is a functional GPCR and has therapeutic potential. Thus our study provides an example that the common noncoding variant with modest effect may provide important biological or therapeutics insights. The strategy we developed here can be applied to other noncoding loci with unknown mechanisms as well.

Our study should be interpreted within the context of its limitations. First, we systematically analyzed all SNPs in 200 Kb window across the locus and found that rs1997243 is the only variant that changes promoter activity of the genome sequence and increases the expression level of *GPR146*. We cannot exclude the possibility that there exist other variants that extremely far away from the lead SNP and mediate the disease-causing effect together with rs1997243. However, our study provides compelling evidences that rs1997243 is the disease-causing variant and increases *GPR146* expression level, which at least contributes to the increased blood cholesterol level in human. Second, our animal models strongly suggest that Gpr146 regulates blood cholesterol level mainly through the liver. However, eQTL analysis showed that the strongest association for *GPR146* expression level and the rs1997243 risk allele is in human whole blood cells. Thus we cannot exclude the possibility that GPR146 may regulate blood cholesterol level through other tissues together with liver in human. Third, we found that *gpr146* knockout mice have decreased blood cholesterol level, however the underline mechanism needs further investigation. Our preliminary data (not shown) indicate that the *gpr146* knockout mice have normal food intake and fecal cholesterol excretion rate, which suggest that the decreased blood cholesterol could be caused by decreased cholesterol secretion into circulation or increased cholesterol clearance from the circulation.

During preparation of this manuscript, Dr. Cowan’s group reported the phenotypic characterization of *gpr*146 knockout mice^23^. They reported that *gpr146* knockout mice have decreased level of blood cholesterol, which is consistent with our results. However, our results provide genetic evidence that GPR146 regulates blood cholesterol level not only in mice but also in human. First, although the 7p22 locus is strongly associated with hypercholesterolemia, we are the first to show that the rs1997243 is the disease-causing variant in this locus. Second, we provide multiple evidences that the rs1997243 risk allele specifically up regulates the expression level of *GPR146* in this locus. Third, by generating the *gpr146* knockout mouse models, we provide strong evidences that Gpr146 positively regulates blood cholesterol level mainly through the liver. Altogether, our results indicate that GPR146 is an important regulator of blood cholesterol level in both human and mouse. Together with the decreased atherosclerosis in *gpr146* knockout mice in Dr. Cowan’s report^23^, we believe GPR146 will be an attractive drug target for hypercholesterolemia and atherosclerotic cardiovascular diseases.

## Methods

### Mice

Mice were housed in the temperature-controlled specific pathogen-free animal facility on a 12 h light-dark daily cycle with free access to water and normal chow diet. All animal care and experimental procedures were approved by the Institutional Animal Use and Care Committee of College of Life Sciences, Wuhan University. *Gpr146 LoxP* mice were generated with CRISPR/Cas9 technology on C57BL/6J background by Nanjing Biomedical Research Institute of Nanjing University. Two gRNA spanning the *gpr146* genome locus are used: 5’-CCAGCAATGCTGGGAGACGT-3’ and 5’-GGCTCCGGGCTCATGTGGGA-3’. Donor vector containing the *gpr146* genome sequence and *LoxP* sites are co-injected with Cas9/gRNA complex (Fig S5a). F0 mice were crossed with wild-type C57BL/6J mice and F1 mice were further genotyped by DNA sequencing, southern blot and PCR analysis. *CMV*-*Cre* (The Jackson Laboratory: 006054), *Album*in*-Cre* (The Jackson Laboratory: 003574) *an*d *Adiponectin*-*Cre* (The Jackson Laboratory: 010803, gift from Dr. Shengzhong Duan, Shanghai Jiao Tong University) mice were used to generate whole body, liver specific and adipose tissue specific *gpr146* knockout mice respectively. High-fat diet (60%, catalog: D12492) and high-fat high-cholesterol diet (40% Cal and 1.25% cholesterol, catalog: D12108C) were obtained from Research diets.

### Cell Culture and Reagents

293T cells, HEK293 cells and HepG2 cells were purchased from China Center for Type Culture Collection (CCTCC) and cultured in high glucose DMEM with 10% FBS and 100 units/mL penicillin G/streptomycin (Gibco, 15140-122). Cells were incubated at 37°C with 5% CO_2_. Cell culture medium was obtained from Life Technologies (12800-082), FBS was obtained from Pan Seratech (ST30-3302). EDTA-free protein inhibitor cocktail was purchased from Bimake (B14001). Polyethylenimine (PEI) was obtained from Polysciences (catalog: 24765, Warrington, PA).

Lentivirus was produced in 293T cells by co-transfecting the packaging plasmids pVSVg (AddGene, 8454) and psPAX2 (AddGene, 12260) with Cas9 (Addgene, 52962) or gRNA (Addgene, 52963) expressing plasmids using PEI according to the instruction. 48 hours after transfection, condition medium was harvest for further experiment.

### CRISPR/dCas9 mediated genome activation

CRISPR/dCas9 mediated genome activation was performed as described^24^ with following modification. The dCas9-GCN4-scFc-p65-HSF1 coding motifs (gift from Dr. Hui Yang, Institute of Neuroscience, Chinese Academy of Sciences) were splitted and expressed separately in pLVX-IRES-Puro vector (Clontech, 632183). pLVX-dCas9-GCN4, PLVX-scFv-p65-HSF1 are packed into lentivirus separately and co-infected with lentivirus expressing gRNA targeting the rs1997243 site in HepG2 cells. 48 hours after infection, cells were collected and RNA was extracted for gene expression analysis.

### Luciferase reporter assay

For promoter activity assay, about 2 kb genome sequence covering indicated SNPs were amplified from HepG2 genome using primers listed in Table S2. The DNA fragments were cloned into upstream of *firefly* luciferase through *Xho1* and *Kpn1* in pGL3-basic vector (Promega, E1751). For enhancer activity assay, a TATA mini promoter was first cloned into upstream of *firefly* luciferase and then DNA fragments were cloned into upstream of TATA box through *Xho1* and *Kpn1* in pGL3-mini promoter vector. Corresponding minor alleles for each SNPs were introduced by recombination with primers listed in Table S2. The *firefly* luciferase reporter plasmids and *renilla* luciferase control plasmid (Promega, E2241) were co-transfected into HepG2 cells with Lipoplus (Sagecreation, Q03003) according to the instruction. 24 hours after transfection, luciferase activity was determined using Dual-luciferase reporter assay system (Promega, E1960). *Firefly* luciferase activity was normalized with *renilla* luciferase activity and the vector transfected group was set to one. Transfection efficiency for each reporters were measured by real-time PCR using DNA extracted from cells in parallel experiments. Specific primers recognizing *firefly* luciferase and *renilla* luciferase are listed in Table S2.

The CRE-luciferase reporter was generated by putting the cAMP regulatory elements (CRE) in front of *firefly* luciferase in pGL4 basic vector (Promega, E134A). *hRenilla* luciferase (Promega, E692A) was used as internal control. To increase plasma membrane localization of GPR146, a 39 amino acids of bovine rhodopsin was fused to the N terminal of GPR146 as described previously^25^. GPR146 expressing plasmid or vector control plasmid was co-transfected with luciferase reporter plasmids by transient transfection using PEI. Eight hours later, cells were changed with serum free medium containing 0.1% fatty acid free BSA (Sangon Biotech, A602448). About 16 hours later, cells were treated with or without indicated amount of fetal bovine serum (FBS) for 6 hours. Luciferase activity was assayed with Dual-luciferase reporter assay system (Promega, E1960). Filtered FBS was made by passing through a 3 kD cut-off filter with centrifugation (Millipore, UFC9003). Heat-inactivated FBS was made by boiling the 3 kD filtered FBS at 95oC for 10 minutes. Then the serum was span down and the supernatant was collected for experiments. In some experiments, cells were pre-incubated with 10 μM of PKA inhibitor H-89 (Selleck, S1582) for 60 minutes before FBS treatment.

### ENCODE analysis

DNase I hypersensitive signal integrates display of DNase I hypersensitivity in multi cell lines by UCSC. DNase I Seq and H3K27ac, H3K4me3 histone modification Chip-seq were performed on human liver samples and the data was downloaded from ENCODE project data portal.

Links for these 4 datasets are:

DNase I hypersensitive signal:

http://genome.ucsc.edu/cgi-bin/hgTrackUi?db=hg38&g=wgEncodeRegDnase

DNase I Seq: https://www.encodeproject.org/experiments/ENCSR158YXM/

H3K27ac Chip-seq: https://www.encodeproject.org/experiments/ENCSR981UJA/

H3K4me3 Chip-seq:https://www.encodeproject.org/experiments/ENCSR344TLI/

### Linkage disequilibrium analysis

Linkage disequilibrium (r^2^) was calculated with Ensemble LD calculator using European population from 1000 genome phase 3. The window size was set to 200 kb around rs1997243. r^2^>0.8 was considered as positive.

### eQTL analysis

The eQTL data in whole blood cells was obtained from gTex and the eQTL figure was generated using gTex webtool.

### Transmembrane domain and variant impact prediction

GPR146 Transmembrane domain was predicted by TMHMM server V.2.0 at http://www.cbs.dtu.dk/services/TMHMM/

*GPR146* p. Gly11Glu impact prediction was performed with two different software.

PolyPhen-2: http://genetics.bwh.harvard.edu/pph2/

PROVEAN: http://provean.jcvi.org/index.php

### Blood chemistry and lipoprotein profiling

Plasma total cholesterol and triglyceride levels were measured by enzymatic kits (Shanghai Kehua Bio-Engineering Co). Plasma lipoprotein particles were size fractionated by Fast Protein Liquid Chromatography (FPLC) using a Superose 6 column (GE Healthcare) and the cholesterol content in each fraction was measured accordingly^26^.

### Immunoblot analysis

Biotinylation and immunoblot was performed as described previously^27^ except that the cells and liver tissue were lysated in RIPA buffer (50 mM Tris-HCl, pH=8.0, 150 mM NaCl, 0.1% SDS, 1.5% NP40, 0.5% deoxycholate, 2 mM MgCl2). For cell fractionation experiments, cells were homogenized by passing through #7 needle for 60 times on ice. Nuclear pellet was isolated by low speed centrifugation (750 g) for 20 minutes at 4°C. The supernatant was transferred for high speed (100,000 g) centrifugation for 60 minutes at 4°C. Then the supernatant was saved as cytosol fraction and the membrane pellet was re-suspended in SDS lysis buffer (10mM Tris-HCl, pH 6.8, 100mM NaCl, 1% SDS, 1mM EDTA, 1 mM EGTA) and incubated at 37 degree for 30 minutes. Then membrane suspension was span down again at 12,000 g for 5 minutes at room temperature and was saved as membrane fraction. The nuclear pellet was re-suspended in nuclear lysate buffer (20mM HEPES/KOH, pH7.6, 2.5% (v/v) glycerol, 1.5mM MgCl_2_, 0.42M NaCl, 1mM EDTA, 1mM EGTA)and rotates at 4 degree for 1 hour. Then the suspension was span down at 14,000 g for 20 minutes at 4oC and the supernatant was saved as nuclear fraction. Cytosol, membrane and nuclear fractions were added with sample buffer and incubated at 37 degree for 30 minutes and then subjected to immunoblot analysis. To detect apolipoproteins, the fresh blood was collected into EDTA coated tubules containing aprotinin (Sigma, A1153-25MG). Then plasma was isolated at 4 degree and was subjected to immunoblots analysis.

The following antibodies were used in this study: anti-Flag antibody (Medical & Biological Laboratories, PM020); anti-calnexin antibody (Proteintech, 10427-2-AP); anti-Actin antibody (Proteintech, 20536-1-AP), anti-Lamin B1 antibody (Proteintech, 20536-1-AP), anti-GPR146 antibody (CUSABIO,CSB-PA006863), anti-ATP1A1 antibody (Abclonal, A0643), anti-ApoB antibody^26^ and anti-ApoA1 antibody (Proteintech, 14427-1-AP).

### Real-Time PCR analysis

Total RNA was extracted from cells or mouse tissues using TRI Reagent (Sigma-Aldrich, T9424). The quality and concentration of RNA were measured with NanoDrop ONE^c^ (Thermo Scientific). cDNA was synthesized from 2 μg of total RNA using a cDNA Reverse Transcription Kit (Thermo Scientific, M1682). The cDNA was quantified by Real-Time PCR using SYBR Green master mix (YEASEN Biotech Co, 11201ES08). Reactions were running in technical duplicate on CFX96 or CFX384 wells plates. Relative quantification was completed using the ΔΔCT method. Gene expression was normalized to housekeeping gene Gapdh, 36B4 or Cyclophilin.

### Primary hepatocytes and non-hepatocytes

Mouse liver was first ligated and a piece of liver was sliced and frozen in liquid N_2_ as whole liver sample. The left over liver was perfused with wash buffer followed by collagenases 1 digestion buffer exactly the same as described^27^. Then the liver was transferred to a 60 mm dish in cold digestion buffer and the particulate material was filtered through a 70 μm filter. The pass through was span down at 40 g*10 minutes for three times at 4°C. The pellets from each spin were pooled together and are the primary hepatocytes. The supernatant was span down again at 500 g for 10 minutes. The pellet was collected as non-hepatocytes. Total RNA was extracted and 2 μg of RNA was subjected to reverse transcription as described above followed by real-time PCR analysis.

### Adeno-associated virus packaging and purification

Adeno-associated virus was produced in 293T cells by co-transfecting the shRNA expression AAV shuttle plasmid together with Delta F6 helper plasmid, Rev Cap 2/9 plasmids using PEI. 60 hours after transfection, cells were harvested and freeze-thaw in lipid nitrogen for five times. Then Benzonase Nuclease (Sigma, E1014) was added and incubated at 37 degree for 45 minutes. After centrifugation at 4000 rpm for 30 minutes, the supernatant was collected and added on top of the iodixanol gradient (15%, 25%, 40%, 58%). Then the sample was centrifuged at 48000 rpm in a Beckman type 70Ti rotor for 130 minutes at 18 degree. The fraction in the 40% iodixanol was collected and dialyzed with 1×PBS extensively. The titer of the virus was determined by qRT-PCR. To knock down *gpr146* in the liver, each mouse was infused with 1X10^11^ virus particles through tail vein and samples were collected at one and two weeks after injection. The shRNA sequence against *gpr146* was listed in Table S2.

### Data analysis

All data are expressed as mean ± SEM and p values were calculated using Student’s test in GraphPad unless otherwise indicated.

## Abbreviations

GWAS: Genome-wide association study
SNPs: Single nucleotide polymorphisms
DHS: DNase I hypersensitive signal
eQTL: Expression quantitative trait loci
HDL: High-density lipoprotein
LDL: Low-density lipoprotein
CRISPR/Cas9: Clustered regularly interspaced short palindromic repeats

## Acknowledgements

The authors would like to thank Dr. Hui Yang (Institute of Neuroscience, Chinese Academy of Sciences) for the dCas9-GCN4-scFc-p65-HSF1 plasmid, the ENCODE Consortium and the ENCODE production laboratory(s) generating the datasets used in the manuscript. This work was supported by grants from the National Natural Science Foundation of China (31570807, 31771304, 91754101, 91857000), the National Key Research and Development Program of China (2016YFA0500100, 2018YFA0800700), the 111 Project of China (B16036), the Fundamental Research Funds for the Central Universities of China (2042017kf0240, 2042017kf0187), the Key Research and Development Program of Hubei province (2019CFA067) and state key laboratory of Natural Medicines (SKLNMKF201902).

## Author contributions

F.F.H, X.L, C.F.C, Y.N.L and Y.W designed the study. F.F.H, X.L, C.F.C, Y.N.L, M.K.D, Y.Y.G, Y.L.Z, X.D.L developed experimental methods. F.F.H, X.L, C.F.C, Y.N.L, M.K.D, Y.Y.G, Y.L.Z, X.D.L, X.M.L performed experiments. D.H.W, Y.Z, M.A, Y.L, B.L.S, H.S.H.H, Y.W contributed to data analysis and interpretation. F.F.H, X.L, C.F.C, Y.N.L and Y.W wrote the manuscript with input from all authors. Y.W supervised the work and obtained the funding.

---

**Figure S1.**
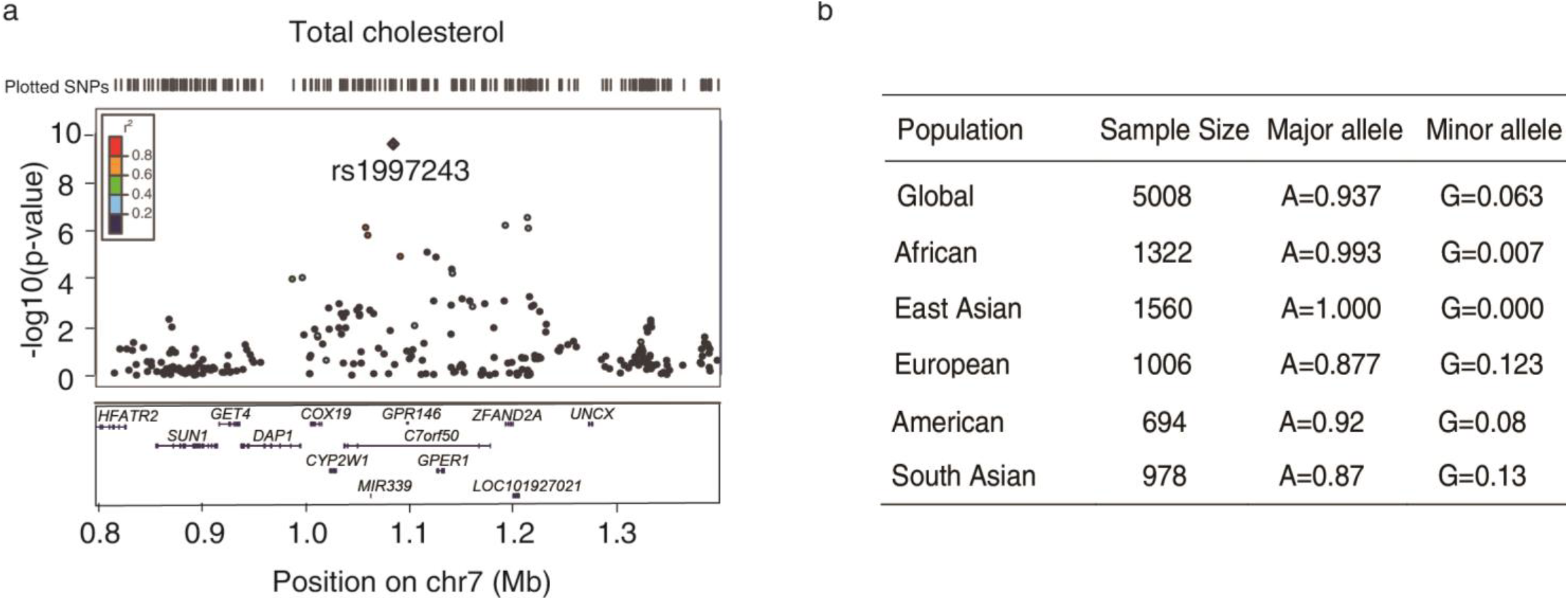
7p22 locus is strongly associated with total cholesterol level. **a**, Reginal (7p22) plot of SNPs associated with total cholesterol level with LocusZoom^14^. Note that rs1997243 is the lead SNP in this locus. **b**, Allele frequency of rs1997243 in different populations from 1000 genome project.

**Figure S2.**
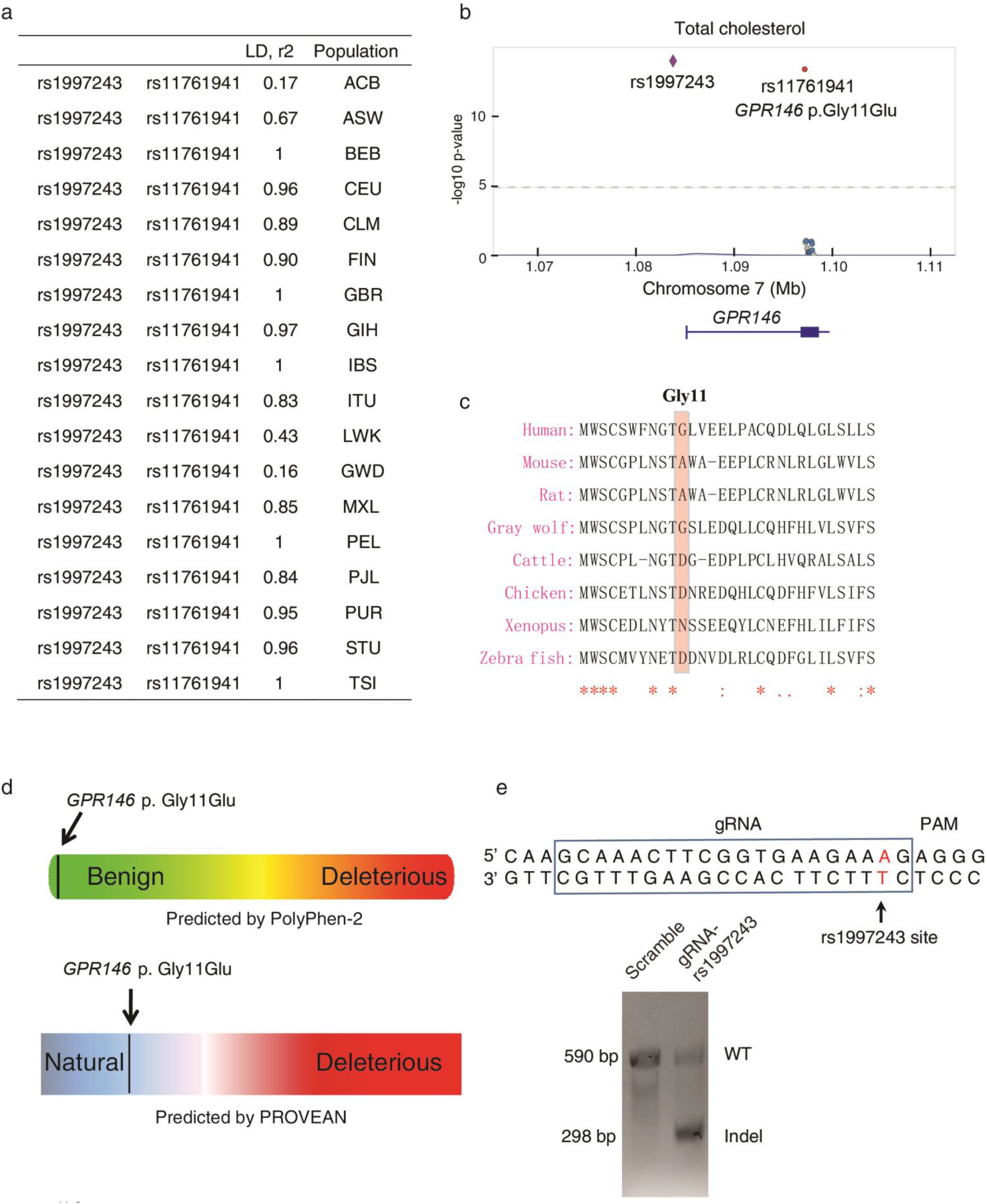
rs11761941 (*GPR146* p. Gly11Glu) has strong linkage disequilibrium with lead SNP rs1997243 and are both associated with blood cholesterol level. **a**, Linkage disequilibrium (r^2^) between rs1997243 and rs11761941 in different populations. LD (r^2^) was calculated with Ensemble LD calculator using data from 1000 genome phase 3. The population codes were exactly the same as described on Ensemble. **b**, Reginal (7p22) plot of SNPs associated with total cholesterol level with LocusZoom^28^. **c**, Sequence alignment of GPR146 across different species. Gly11 is not conserved and has been substituted with Asp, Asn or Ala in other species except *Gray wolf*. **d**, *GPR146* p. Gly11Glu is predicted to be benign. Different software all predicts that the *GPR146* p. Gly11Glu variant is benign and natural. **e**, gRNA sequence targeting rs1997243 site. Boxed sequence is the gRNA sequence followed by the PAM sequence AGG. HepG2 cells stably expressing wild-type spCas9 were infected with lentivirus expressing this gRNA. The gRNA efficiency was evaluated with T7 endonuclease I digestion as described in Methods (lower panel). The result indicates that this gRNA is effective in targeting rs1997243 site and is used for enzymatic dead Cas9-mediated activation system (Fig 1e).

**Figure S3.**
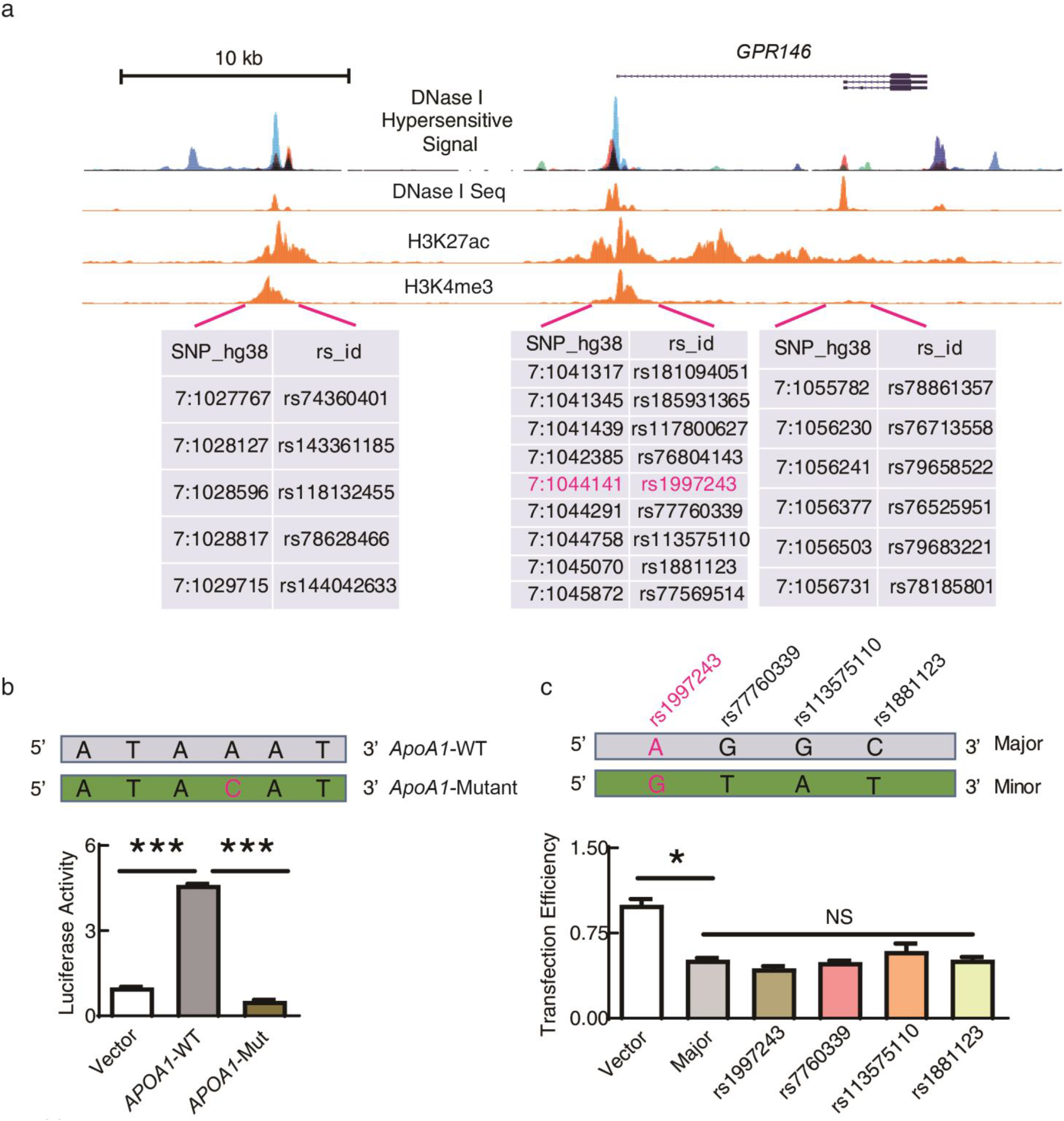
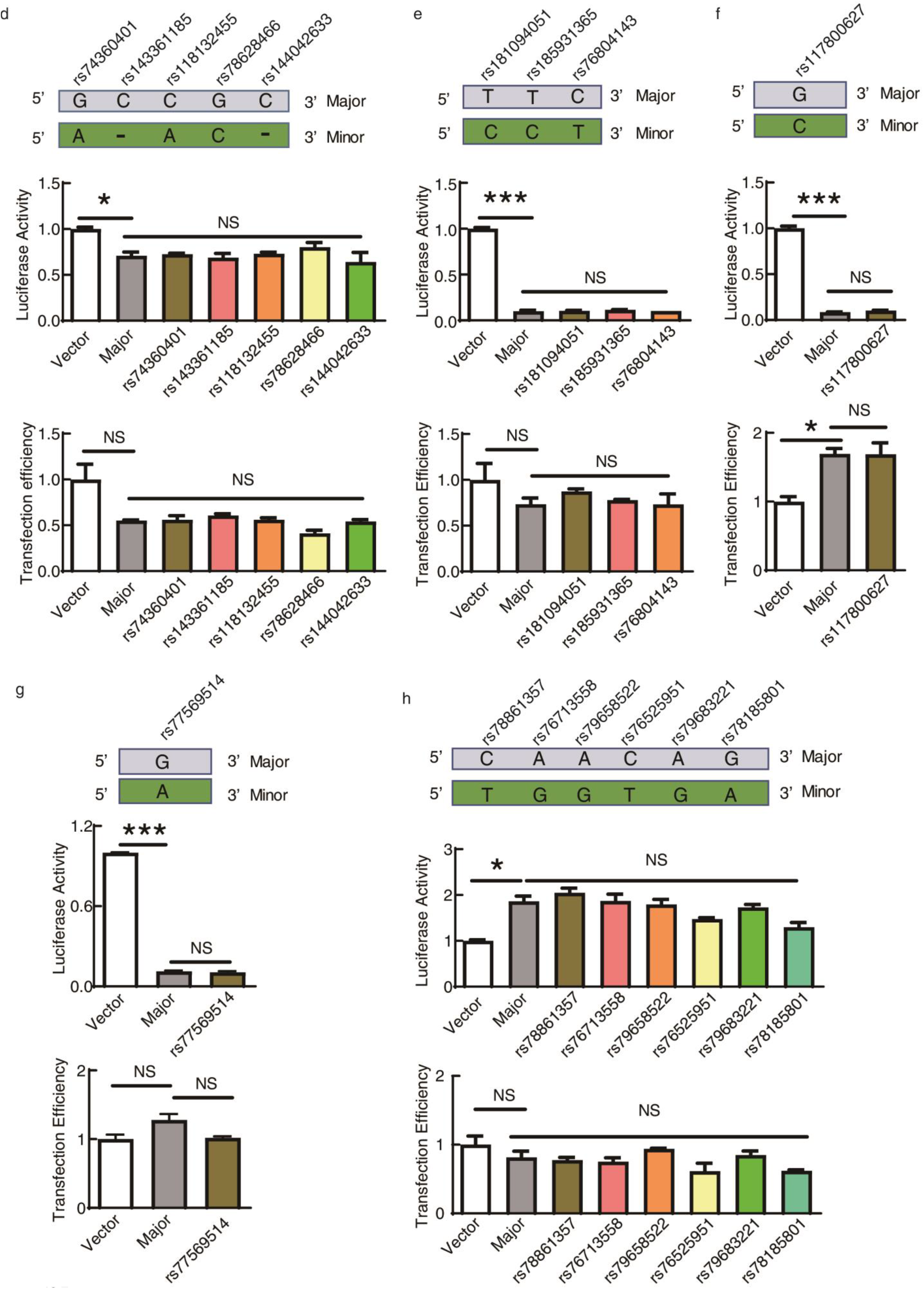
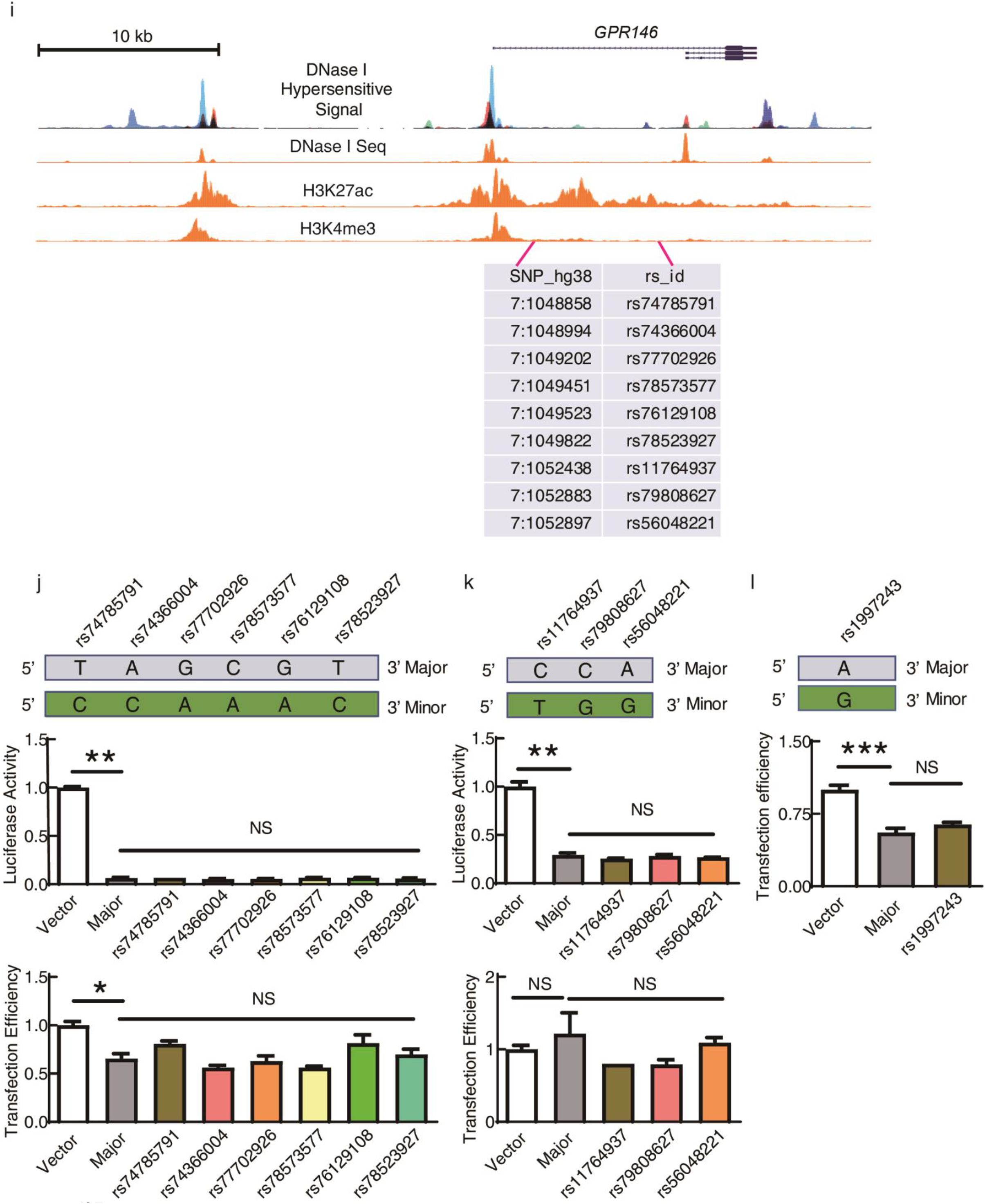
Characterization of SNPs in 7p22 locus. **a,** SNPs that have strong linkage disequilibrium with lead SNP rs1997243 and are located in genome active regions are listed in the tables. Linkage disequilibrium (LD, r^2^) was calculated with data from 1000 genome phase 3 as described in Methods. Genome active regions were identified by DNase I hypersensitive signal from 95 cell lines and DNase I seq, H3K27ac Chip-seq, H3K4me3 Chip-seq signals from human liver samples. SNPs with r^2^ >0.8 and localized in the genome active regions are listed in the tables and subjected to further functional study. **b**, Luciferase reporter activity for *ApoA1* promoter sequence which is used as control for the assay. *ApoA1* wild-type (WT) promoter or its single base pair mutant was cloned into upstream of *firefly* luciferase. *Firefly* luciferase activity was measured and normalized with *renilla* luciferase activity, with vector group was set to 1. **c**, Transfection efficiency for experiment described in Fig 1b. After transfection, cells were harvested and genomes DNA were extracted and subjected to real-time PCR analyze as described in Methods. **d-h**, Luciferase reporter activities and their transfection efficiency for all SNPs listed in Fig S3a as described in Fig 1b and Fig S3c. **i**, SNPs have strong linkage disequilibrium with lead SNP rs1997243 and are located in H3K27ac markered region. **j-k**, Luciferase reporter activities and their transfection efficiency for all SNPs listed in Fig S3i. **l**, Transfection efficiency for experiment Fig 1c. All data are expressed as means ± S EM and *p* values were calculated using Student’s test (\**p*<0.05, \*\**p*<0.01, \*\*\**p*<0.001). All experiments were repeated at with similar results.

**Figure S4.**
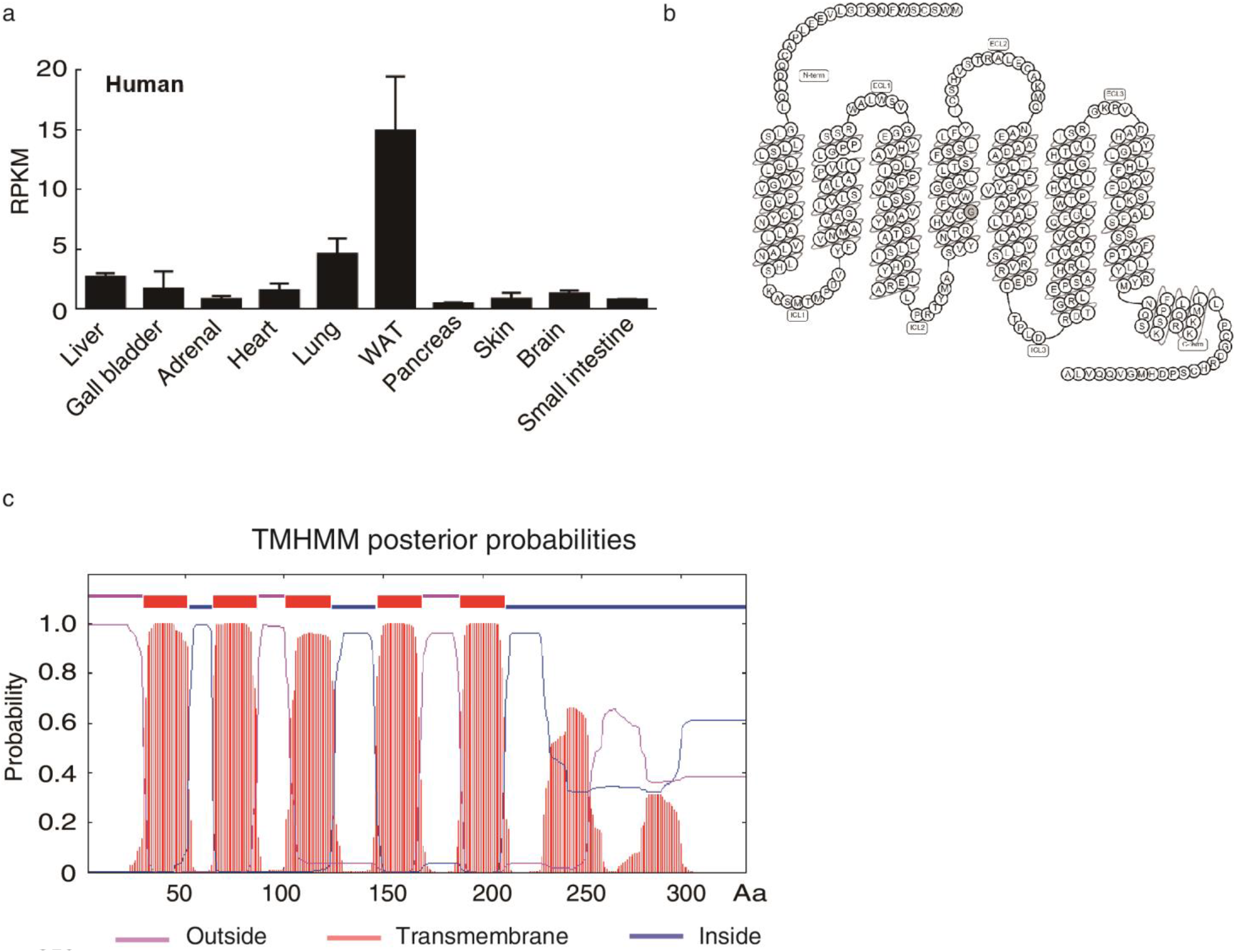
Gpr146 is a typical GPCR that is specifically expressed in hepatocytes of mouse liver. **a**, Tissue distribution of *GPR146* in normal human tissues. Human *GPR146* expression data was downloaded from NCBI with HPA RNA-seq normal tissues dataset. **b**, Topology of human GPR146. The human GPR146 topology was generated from gpcr database (https://www.gpcrdb.org/). **d**, Human GPR146 is predicted to have 7 transmembrane domains with N terminal facing extracellular compartment. Transmembrane domain prediction was performed with TMHMM server V.2.0 as described in Methods.

**Figure S5.**
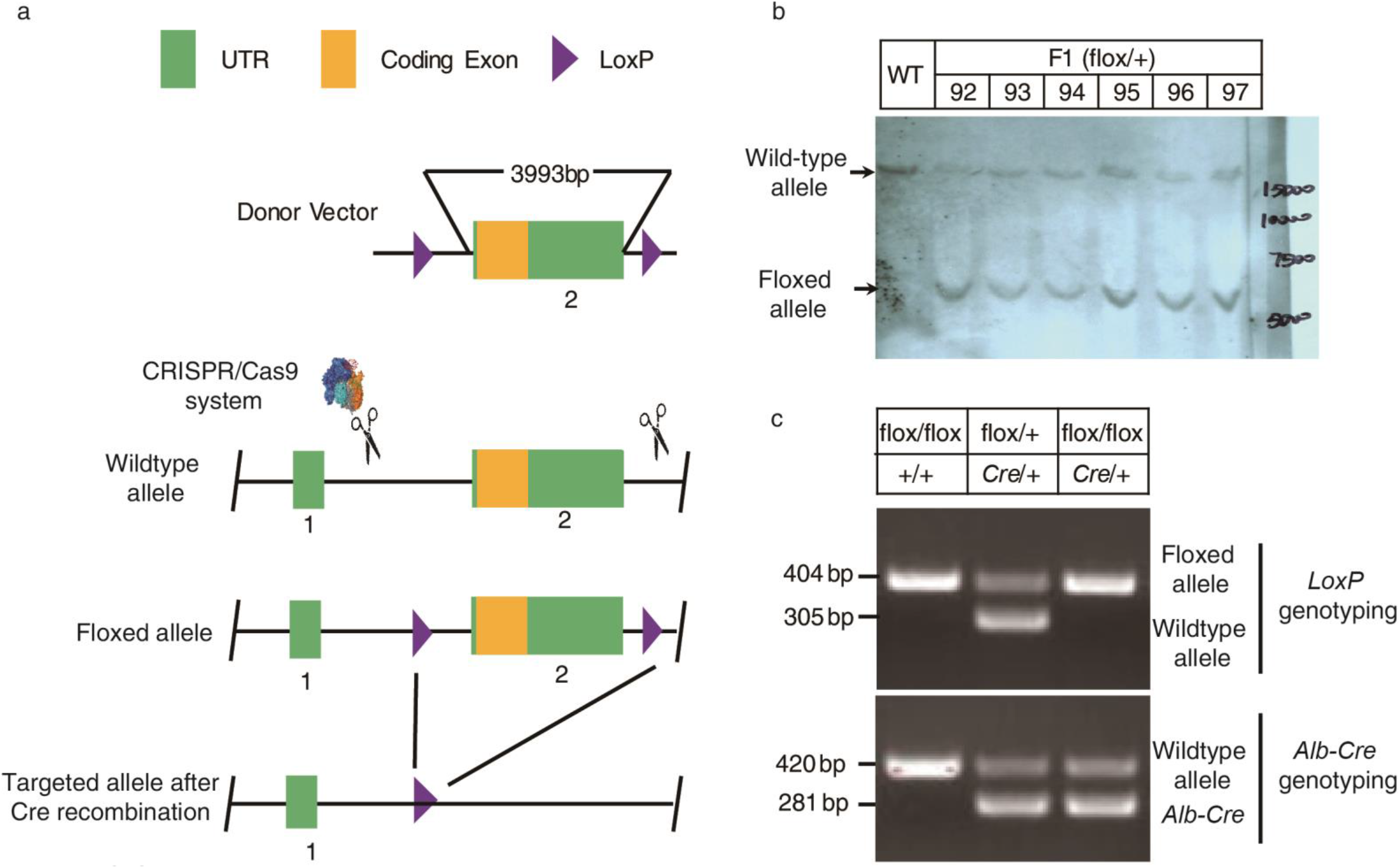
Generating *gpr146* conditional knockout mice with Cre-LoxP system. **a**, Schematic diagram showing the generation of *gpr146 LoxP* mice with CRISPR/Cas9 system as described in Methods. **b**, Southern blot verification of the *LoxP* allele. Genome DNA was extracted from mouse tail and subjected to southern blot analysis. Six F1 heterozygous mice were genotyped and wild-type (WT) mice were used as control. **c**, PCR genotyping of heterozygous and homozygous liver specific *gpr146* knockout mice.

**Figure S6.**
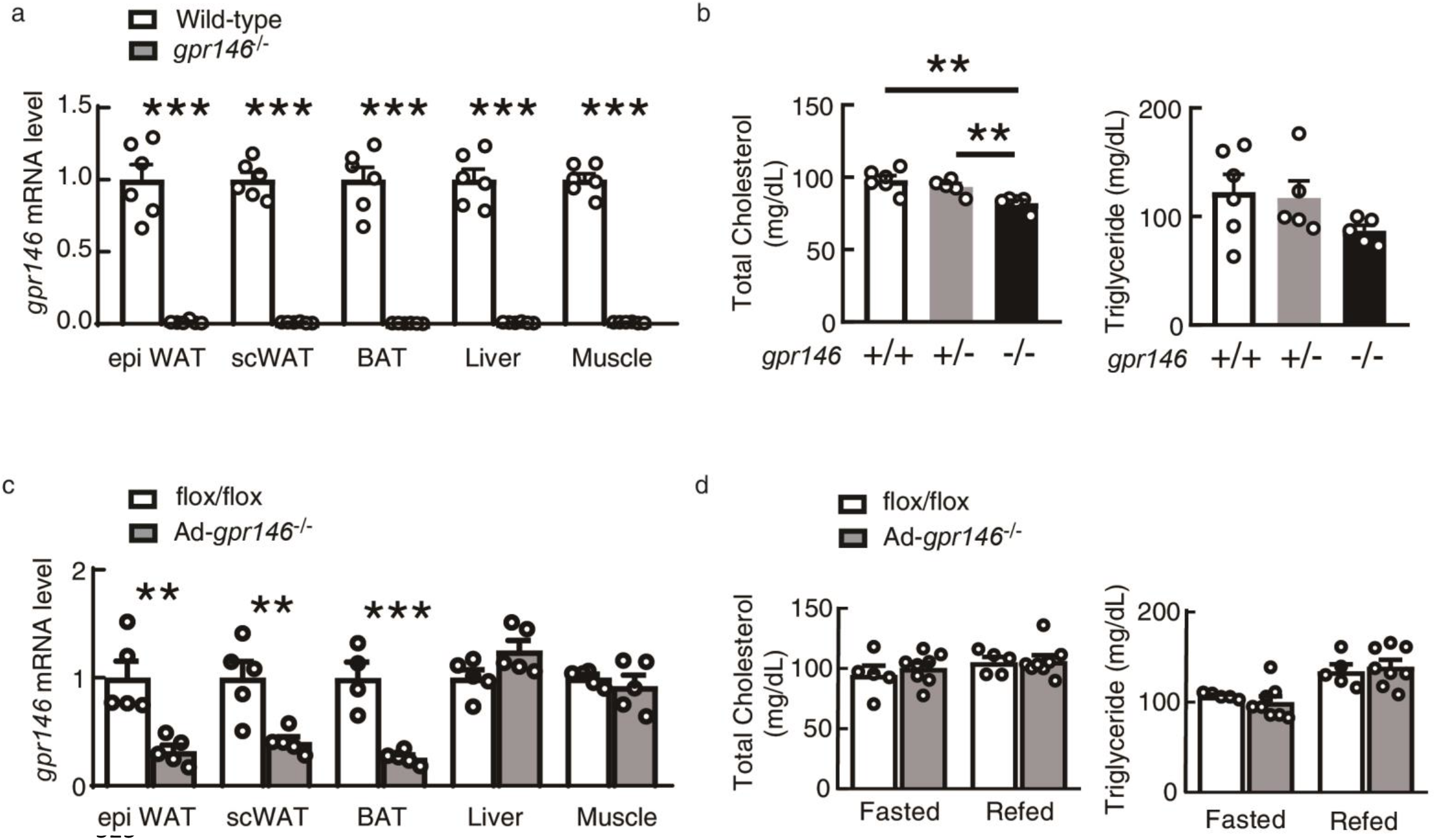
Phenotypic characterization of whole body and adipose tissue specific *gpr146* knockout mice. **a**, mRNA levels of *Gpr146* in tissues of whole body *gpr146* knockout mice (*gpr146*^−/−^) and their littermate controls (n=6/group, female, 12-15 weeks). **b**, Plasma levels of total cholesterol and triglyceride in overnight fasted heterozygous, homozygous whole body *gpr146* knockout mice and their littermate controls (n=5-6/group, male, 8-10 weeks). **c**, *Gpr146* mRNA levels in tissues of adipose tissue specific *gpr146* knockout mice (Ad-*gpr146*^−/−^) and their littermate controls (n=4-5/group, male, 9-10 weeks). **d**, Plasma levels of total cholesterol and triglyceride in Ad-*gpr146*^−/−^ mice and their littermate controls at 16 hours fasting or 6 hours refeeding after a 16 hours fasting (n=5-8/group, male, 9-10 weeks). epiWAT, epididymal white adipose tissue; scWAT, subcutaneous white adipose tissue; BAT, brown adipose tissue. All data are expressed as means ± SEM and *p* values were calculated using Student’s test (\*\**p*<0.01, \*\*\**p*<0.001). All experiments were repeated with similar results.

## Reference

1. Hindorff, L.A.e.a. A Catalog of Published Genome-Wide Association Studies. Available at www.genome.gov/gwastudies.

2. Maurano, M.T. et al. Systematic localization of common disease-associated variation in regulatory DNA. Science 337, 1190–5 (2012).

3. Kathiresan, S. et al. A genome-wide association study for blood lipid phenotypes in the Framingham Heart Study. BMC Med Genet 8 Suppl 1, S17 (2007).

4. Weiss, L.A., Pan, L., Abney, M. & Ober, C. The sex-specific genetic architecture of quantitative traits in humans. Nat Genet 38, 218–22 (2006).

5. Chen, L. et al. Regulation of glucose and lipid metabolism in health and disease. Sci China Life Sci 62, 1420–1458 (2019).

6. Goldstein, J.L. & Brown, M.S. The LDL receptor. Arterioscler Thromb Vasc Biol 29, 431–8 (2009).

7. Garcia, C.K. et al. Autosomal recessive hypercholesterolemia caused by mutations in a putative LDL receptor adaptor protein. Science 292, 1394–8 (2001).

8. Brooks-Wilson, A. et al. Mutations in ABC1 in Tangier disease and familial high-density lipoprotein deficiency. Nat Genet 22, 336–45 (1999).

9. Berge, K.E. et al. Accumulation of dietary cholesterol in sitosterolemia caused by mutations in adjacent ABC transporters. Science 290, 1771–5 (2000).

10. Abifadel, M. et al. Mutations in PCSK9 cause autosomal dominant hypercholesterolemia. Nat Genet 34, 154–6 (2003).

11. Altmann, S.W. et al. Niemann-Pick C1 Like 1 protein is critical for intestinal cholesterol absorption. Science 303, 1201–4 (2004).

12. Wang, L.J. et al. Molecular characterization of the NPC1L1 variants identified from cholesterol low absorbers. J Biol Chem 286, 7397–408 (2011).

13. Zhang, Y.Y. et al. A LIMA1 variant promotes low plasma LDL cholesterol and decreases intestinal cholesterol absorption. Science 360, 1087–1092 (2018).

14. Willer, C.J. et al. Discovery and refinement of loci associated with lipid levels. Nat Genet 45, 1274–1283 (2013).

15. Liu, D.J. et al. Exome-wide association study of plasma lipids in >300,000 individuals. Nat Genet 49, 1758–1766 (2017).

16. Klarin, D. et al. Genetics of blood lipids among ~300,000 multi-ethnic participants of the Million Veteran Program. Nat Genet 50, 1514–1523 (2018).

17. Consortium, E.P. An integrated encyclopedia of DNA elements in the human genome. Nature 489, 57–74 (2012).

18. Gross, D.S. & Garrard, W.T. Nuclease hypersensitive sites in chromatin. Annu Rev Biochem 57, 159–97 (1988).

19. Thurman, R.E. et al. The accessible chromatin landscape of the human genome. Nature 489, 75–82 (2012).

20. Heintzman, N.D. et al. Distinct and predictive chromatin signatures of transcriptional promoters and enhancers in the human genome. Nat Genet 39, 311–8 (2007).

21. Ernst, J. & Kellis, M. Discovery and characterization of chromatin states for systematic annotation of the human genome. Nat Biotechnol 28, 817–25 (2010).

22. Cheng, Z. et al. Luciferase Reporter Assay System for Deciphering GPCR Pathways. Curr Chem Genomics 4, 84–91 (2010).

23. Yu, H. et al. GPR146 Deficiency Protects against Hypercholesterolemia and Atherosclerosis. Cell 179, 1276–1288 e14 (2019).

24. Zhou, H. et al. In vivo simultaneous transcriptional activation of multiple genes in the brain using CRISPR-dCas9-activator transgenic mice. Nat Neurosci 21, 440–446 (2018).

25. Chandrashekar, J. et al. T2Rs function as bitter taste receptors. Cell 100, 703–11 (2000).

26. Wang, Y. et al. Inactivation of ANGPTL3 reduces hepatic VLDL-triglyceride secretion. J Lipid Res 56, 1296–307 (2015).

27. Wang, Y., Huang, Y., Hobbs, H.H. & Cohen, J.C. Molecular characterization of proprotein convertase subtilisin/kexin type 9-mediated degradation of the LDLR. J Lipid Res 53, 1932–43 (2012).

28. Lu, X. et al. Exome chip meta-analysis identifies novel loci and East Asian-specific coding variants that contribute to lipid levels and coronary artery disease. Nat Genet 49, 1722–1730 (2017).

