## Supplementary Tables for "Hypercholesterolemia risk associated GPR146 is an orphan G-protein coupled receptor that regulates blood cholesterol level in human and mouse"

**Supplementary table 1**, SNPs that have strong linkage disequilibrium with lead SNP rs1997243. The linkage disequilibrium (LD, r2) was calculated using European population datasets from 1000 genome phase 3 as described in Methods. LD with r2>0.8 are listed and SNPs located in genome active regions are highlighted and subjected to further function studies.

| Variant 1 | Variant 1 location | Variant 2 | Variant 2 location | LD ($r^2$) |
|---|---|---|---|---|
| rs1997243 | 7:1044141 | rs6951245 | 7:1018557 | 1 |
| rs1997243 | 7:1044141 | rs75280240 | 7:1018875 | 1 |
| rs1997243 | 7:1044141 | rs57068908 | 7:1020448 | 1 |
| rs1997243 | 7:1044141 | rs11763020 | 7:1020652 | 1 |
| rs1997243 | 7:1044141 | rs11763765 | 7:1020874 | 1 |
| rs1997243 | 7:1044141 | rs1881128 | 7:1023677 | 1 |
| rs1997243 | 7:1044141 | rs11766669 | 7:1023957 | 1 |
| rs1997243 | 7:1044141 | rs11768895 | 7:1024055 | 0.803571 |
| rs1997243 | 7:1044141 | rs79067319 | 7:1025912 | 1 |
| rs1997243 | 7:1044141 | rs884977 | 7:1026774 | 1 |
| rs1997243 | 7:1044141 | rs74360401 | 7:1027767 | 1 |
| rs1997243 | 7:1044141 | rs143361185 | 7:1028127 | 1 |
| rs1997243 | 7:1044141 | rs118132455 | 7:1028596 | 1 |
| rs1997243 | 7:1044141 | rs78628466 | 7:1028817 | 1 |
| rs1997243 | 7:1044141 | rs144042633 | 7:1029715 | 1 |
| rs1997243 | 7:1044141 | rs11768761 | 7:1030171 | 1 |
| rs1997243 | 7:1044141 | rs76833820 | 7:1030642 | 1 |
| rs1997243 | 7:1044141 | rs28379681 | 7:1030903 | 1 |
| rs1997243 | 7:1044141 | rs28399710 | 7:1030995 | 1 |
| rs1997243 | 7:1044141 | rs28528096 | 7:1031276 | 1 |
| rs1997243 | 7:1044141 | rs112425403 | 7:1031895 | 0.961981 |
| rs1997243 | 7:1044141 | rs74369356 | 7:1032075 | 1 |
| rs1997243 | 7:1044141 | rs77083167 | 7:1032187 | 1 |
| rs1997243 | 7:1044141 | rs199718264 | 7:1032483 | 1 |
| rs1997243 | 7:1044141 | rs74347384 | 7:1032804 | 1 |
| rs1997243 | 7:1044141 | rs78308415 | 7:1032998 | 1 |
| rs1997243 | 7:1044141 | rs75493422 | 7:1033254 | 1 |
| rs1997243 | 7:1044141 | rs118059236 | 7:1033566 | 1 |
| rs1997243 | 7:1044141 | rs113858334 | 7:1033840 | 1 |
| rs1997243 | 7:1044141 | rs79765398 | 7:1034128 | 1 |
| rs1997243 | 7:1044141 | rs79443843 | 7:1035609 | 1 |
| rs1997243 | 7:1044141 | rs113713890 | 7:1036436 | 1 |
| rs1997243 | 7:1044141 | rs78896566 | 7:1040546 | 1 |
| rs1997243 | 7:1044141 | rs79422648 | 7:1040605 | 1 |
| rs1997243 | 7:1044141 | rs113119264 | 7:1040928 | 1 |
| rs1997243 | 7:1044141 | rs140138013 | 7:1041233 | 0.960947 |
| rs1997243 | 7:1044141 | rs186044114 | 7:1041261 | 0.922562 |
| rs1997243 | 7:1044141 | rs145122769 | 7:1041289 | 0.960947 |
| rs1997243 | 7:1044141 | rs181094051 | 7:1041317 | 0.960947 |
| rs1997243 | 7:1044141 | rs185931365 | 7:1041345 | 1 |
| rs1997243 | 7:1044141 | rs117800627 | 7:1041439 | 0.852872 |
| rs1997243 | 7:1044141 | rs76804143 | 7:1042385 | 1 |
| rs1997243 | 7:1044141 | rs77760339 | 7:1044291 | 1 |
| rs1997243 | 7:1044141 | rs113575110 | 7:1044758 | 1 |
| rs1997243 | 7:1044141 | rs1881123 | 7:1045070 | 1 |
| rs1997243 | 7:1044141 | rs77569514 | 7:1045872 | 1 |

| | | | | |
|---|---|---|---|---|
| rs1997243 | 7:1044141 | rs148758091 | 7:1046716 | 1 |
| rs1997243 | 7:1044141 | rs11770909 | 7:1047080 | 1 |
| rs1997243 | 7:1044141 | rs74785791 | 7:1048858 | 1 |
| rs1997243 | 7:1044141 | rs74366004 | 7:1048994 | 1 |
| rs1997243 | 7:1044141 | rs77702926 | 7:1049202 | 1 |
| rs1997243 | 7:1044141 | rs78573577 | 7:1049451 | 1 |
| rs1997243 | 7:1044141 | rs76129108 | 7:1049523 | 1 |
| rs1997243 | 7:1044141 | rs78523927 | 7:1049822 | 1 |
| rs1997243 | 7:1044141 | rs74652290 | 7:1050473 | 1 |
| rs1997243 | 7:1044141 | rs78185558 | 7:1050547 | 1 |
| rs1997243 | 7:1044141 | rs78158942 | 7:1050804 | 1 |
| rs1997243 | 7:1044141 | rs2363286 | 7:1051989 | 1 |
| rs1997243 | 7:1044141 | rs11764937 | 7:1052438 | 0.926339 |
| rs1997243 | 7:1044141 | rs79808627 | 7:1052883 | 0.860168 |
| rs1997243 | 7:1044141 | rs56048221 | 7:1052897 | 0.821014 |
| rs1997243 | 7:1044141 | rs77868187 | 7:1054332 | 0.961981 |
| rs1997243 | 7:1044141 | rs75016635 | 7:1054485 | 0.961981 |
| rs1997243 | 7:1044141 | rs113066613 | 7:1054492 | 0.961981 |
| rs1997243 | 7:1044141 | rs11763793 | 7:1054706 | 0.961981 |
| rs1997243 | 7:1044141 | rs11764748 | 7:1054872 | 0.961981 |
| rs1997243 | 7:1044141 | rs11763835 | 7:1054877 | 0.961981 |
| rs1997243 | 7:1044141 | rs78861357 | 7:1055782 | 0.961981 |
| rs1997243 | 7:1044141 | rs76713558 | 7:1056230 | 0.961981 |
| rs1997243 | 7:1044141 | rs79658522 | 7:1056241 | 0.961981 |
| rs1997243 | 7:1044141 | rs76525951 | 7:1056377 | 0.961981 |
| rs1997243 | 7:1044141 | rs79683221 | 7:1056503 | 0.961981 |
| rs1997243 | 7:1044141 | rs78185801 | 7:1056731 | 0.961981 |
| rs1997243 | 7:1044141 | rs80031817 | 7:1057157 | 0.961981 |
| rs1997243 | 7:1044141 | rs78143408 | 7:1057210 | 0.961981 |
| rs1997243 | 7:1044141 | rs78351779 | 7:1057387 | 0.961981 |
| rs1997243 | 7:1044141 | rs11761941 | 7:1057547 | 0.961981 |
| rs1997243 | 7:1044141 | rs11767527 | 7:1057758 | 0.961981 |
| rs1997243 | 7:1044141 | rs61910751 | 7:1058259 | 0.961981 |
| rs1997243 | 7:1044141 | rs77434655 | 7:1058769 | 0.961981 |
| rs1997243 | 7:1044141 | rs75398423 | 7:1058778 | 0.961981 |
| rs1997243 | 7:1044141 | rs112309216 | 7:1060660 | 0.961981 |
| rs1997243 | 7:1044141 | rs76161580 | 7:1060671 | 0.961981 |
| rs1997243 | 7:1044141 | rs76388414 | 7:1061058 | 0.961981 |
| rs1997243 | 7:1044141 | rs77305932 | 7:1061225 | 0.961981 |
| rs1997243 | 7:1044141 | rs79788515 | 7:1061443 | 0.926339 |
| rs1997243 | 7:1044141 | rs11768486 | 7:1061654 | 0.961981 |
| rs1997243 | 7:1044141 | rs11766526 | 7:1061797 | 0.961981 |
| rs1997243 | 7:1044141 | rs76214082 | 7:1062461 | 0.961981 |
| rs1997243 | 7:1044141 | rs75488469 | 7:1062486 | 0.961981 |
| rs1997243 | 7:1044141 | rs80094748 | 7:1064322 | 0.961981 |
| rs1997243 | 7:1044141 | rs113146460 | 7:1065528 | 0.961981 |
| rs1997243 | 7:1044141 | rs61753396 | 7:1065736 | 0.961981 |
| rs1997243 | 7:1044141 | rs113642700 | 7:1066018 | 0.961981 |
| rs1997243 | 7:1044141 | rs149125341 | 7:1066258 | 0.961981 |
| rs1997243 | 7:1044141 | rs553221139 | 7:1066867 | 0.961981 |
| rs1997243 | 7:1044141 | rs111705570 | 7:1067106 | 0.961981 |
| rs1997243 | 7:1044141 | rs79327308 | 7:1068505 | 0.961981 |
| rs1997243 | 7:1044141 | rs117729148 | 7:1068895 | 0.961981 |
| rs1997243 | 7:1044141 | rs113221697 | 7:1069107 | 0.961981 |

| | | | | |
|---|---|---|---|---|
| rs1997243 | 7:1044141 | rs113365567 | 7:1069149 | 0.961981 |
| rs1997243 | 7:1044141 | rs113448118 | 7:1069318 | 0.961981 |
| rs1997243 | 7:1044141 | rs79143504 | 7:1070576 | 0.961981 |
| rs1997243 | 7:1044141 | rs574980713 | 7:1070621 | 0.961981 |
| rs1997243 | 7:1044141 | rs111161354 | 7:1070686 | 0.961981 |
| rs1997243 | 7:1044141 | rs532319453 | 7:1070844 | 0.961981 |
| rs1997243 | 7:1044141 | rs558665380 | 7:1070854 | 0.961981 |
| rs1997243 | 7:1044141 | rs190638559 | 7:1070869 | 0.961981 |
| rs1997243 | 7:1044141 | rs200416508 | 7:1070909 | 0.961981 |
| rs1997243 | 7:1044141 | rs74887741 | 7:1071022 | 0.961981 |
| rs1997243 | 7:1044141 | rs79617366 | 7:1072322 | 0.961981 |
| rs1997243 | 7:1044141 | rs111899361 | 7:1072712 | 0.961981 |
| rs1997243 | 7:1044141 | rs80212261 | 7:1074376 | 0.961981 |
| rs1997243 | 7:1044141 | rs77346188 | 7:1074387 | 0.961981 |
| rs1997243 | 7:1044141 | rs77943789 | 7:1100522 | 0.831632 |
| rs1997243 | 7:1044141 | rs10266519 | 7:1103341 | 0.86134 |
| rs1997243 | 7:1044141 | rs74976697 | 7:1120869 | 0.803571 |
| rs1997243 | 7:1044141 | rs28600085 | 7:1130000 | 0.803571 |
| rs1997243 | 7:1044141 | rs28483034 | 7:1130535 | 0.803571 |
| rs1997243 | 7:1044141 | rs183256883 | 7:1135851 | 0.803571 |
| rs1997243 | 7:1044141 | rs6952546 | 7:948373 | 0.883578 |
| rs1997243 | 7:1044141 | rs113184427 | 7:950403 | 0.922546 |
| rs1997243 | 7:1044141 | rs113766238 | 7:950606 | 0.922546 |
| rs1997243 | 7:1044141 | rs79672483 | 7:955120 | 0.922546 |
| rs1997243 | 7:1044141 | rs113464035 | 7:955606 | 0.922546 |

Note: LD data were generated on Ensemble using 1000 genome phase 3 data, with European population. The windown is 200kb. Highlighted SNPs are the one localized in genomic active regions and used for function studies.

**Supplementary Table 2: This table contains sequence information used in this study.**

**shRNA sequence for mouse *gpr146*.**

| Target Gene | Sequence |
|---|---|
| *Gpr146* | GCTTAGGCTATAATGCTCTTC |

**gRNA sequence for rs1997243 site**

| Target Site | Sequence |
|---|---|
| rs1997243 | GCAAACTTCGGTGAAGAAAG |

**Primers for qRT-PCR.**

| **Human** | Forward | Reverse |
|---|---|---|
| *CYP2W1* | CCGATTTGACTACCGGGACC | CGTCCACATAGCTGCACAC |
| *GPER1* | CACCAGCAGTACGTGATCGG | CATCTTCTCGCGGAAGCTGAT |
| *C7orf50* | AGCTGAAAAAGGAACGGAAGAA | TGGAGAAGTGCTCATCGGGAA |
| *COX19* | TTTCGGGACCAAGAGCTTCC | GCATCAATTTTCTCTCCATCCTGC |
| *ZFAND2A* | TGTGACTCTCACCCTGGGAA | GGGTGAGCACCCAGCTT |
| *GPR146* | TGAGCCTCGACCACTACATC | GCTTCTGCGTTCTGCATCTTG |
| *36B4* | TGCATCAGTACCCCATTCTATCA | AAGGTGTAATCCGTCTCCACAGA |
| **Mouse** | Forward | Reverse |
| *Albumin* | CGAGAAGCTTGGAGAATATGGA | CTTGGTGCCCACTCTTCCTA |
| *Apob* | CGTGGGCTCCAGCATTCTA | TCACCAGTCATTTCTGCCTTTG |
| *F4/80* | CTTTGGCTATGGGCTTCCAGTC | GCAAGGAGGACAGAGTTTATCGTG |
| *Gpr146* | TGACCATGTACTCCACTGCAC | AAGACACGTGACTGCAGATGT |
| *36b4* | CACTGGTCTAGGACCCGAGAAG | GGTGCCTCTGGAGATTTTCG |
| *Gapdh* | TGTGTCCGTCGTGGATCTGA | CCTGCTTCACCACCTTCTTGAT |
| *Cyclophilin* | TGGAGAGCACCAAGACAGACA | TGCCGGAGTCGACAATGAT |

**Primers used to amplify genome sequence from HepG2 cells and primers used to introduce mutations for each SNPs.**

| Figures | Primer sequence | |
|---|---|---|
| Fig 1**b-c** | **Reference allele primer** | |
| | F:ATTTCTCTATCGATAGGTACCATGGTGACCTATCTCATTGACGC | |
| | R:ACTTAGATCGCAGATCTCGAGGAATGGAGGACTCAAGTACATCTTCA | |
| | **Primers for mutagenesis** | |
| | rs1997243 | F:AAACTTCGGTGAAGAAGGAGGGGCAGGTGT |
| | | R:CTTCTTCACCGAAGTTTGCTTGCTGTTTCCA |
| | rs77760339 | F:CCGCGCTGGGCACTAAGATAATTCTCAA |
| | | R:ATCTTAGTGCCCAGCGCGGTTTC |
| | rs113575110 | F:TCCCACGGCGGACGCTGCGCTTACTG |
| | | R:CAGCGTCCGCCGTGGGAGCCACAGCAGCTT |
| | rs1881123 | F:TGGATCTGCTGTCAGTTGTCTTGGACAGTT |
| | | R:ACAACTGACAGCAGATCCTACACCTCCATGTA |

| | |
|---|---|
| Fig S3b | **Reference allele primer**<br>F:ATTTCTCTATCGATAGGTACCCTCTGCCAACACAATGGAC<br>R:ACTTAGATCGCAGATCTCGAGGAGCGGGAGAAGACCTCAGG<br>**Primers for mutagenesis**<br>F:ACATACATAGGCCCTGCAAGAGC<br>R:GCCTATGTATGTCTGCAGCCAGG |
| Fig S3d | **Reference allele primer**<br>F:ATTTCTCTATCGATAGGTACCGCCACCGCCTCCTCAGAA<br>R:ACTTAGATCGCAGATCTCGAGCTGTGTGGGCTGGCGCAT |
| | **Primers for mutagenesis**<br>rs74360401 F:CGGCACAGGGAGAGGCGAGGCGGCG<br>R:CGCCGCCTCGCCTCTCCCTGTGCCG<br>rs143361185 F:CGGAGCGGGGCCGCAGCAGCGTCTG<br>R:CAGACGCTGCTGCGGCCCCGCTCCG<br>rs118132455 F:GAACTGCGGGGAGGGCGAACCCCCGCCCGCGCAG<br>R:CTGCGCGGGCGGGGGTTCGCCCTCCCCGCAGTTC<br>rs78628466 F:CAATCTGACTCCATGCCCCCCATCTCCTCC<br>R:GGAGGAGATGGGGGGCATGGAGTCAGATTG<br>rs144042633 F:GGTCGCGGGGCCTGCTCTGGAGGCC<br>R:GGCCTCCAGAGCAGGCCCCGCGACC |
| Fig S3e | **Reference allele primer**<br>F:ATTTCTCTATCGATAGGTACCGGAGGATGAGCAAAGGCAGA<br>R:ACTTAGATCGCAGATCTCGAGCCACGCCTGCTGATGTGATA |
| | **Primers for mutagenesis**<br>rs181094051 Minor allele sequence was synthesized due to multi-repeats near this SNP<br>rs185931365 Minor allele sequence was synthesized due to multi-repeats near this SNP<br>rs76804143 F:TCCAGCTCCTGACCAGCA<br>R:AGATTGGGGCGTAGGGGGGCATGTGGCCCTGACTCAGC |
| Fig S3f | **Reference allele primer**<br>F:ATTTCTCTATCGATAGGTACCAATAGGACTTGGGCTCCGCA<br>R:ACTTAGATCGCAGATCTCGAGAGACAGCATGTCATCATCTGACACT |
| | rs117800627 Minor allele sequence was synthesized directly due to multi-repeats near this SNP. The sequence was further confirmed by sanger sequencing. |
| Fig S3g | **Reference allele primer**<br>F:ATTTCTCTATCGATAGGTACCGTTAGTATTACAAACCTGGCAAGTCAG<br>R:ACTTAGATCGCAGATCTCGAGGGATTCCAGGTGACATCCTAGG |
| | **Primers for mutagenesis**<br>rs77569514 F:ACACATCCGTGCTGGCATCCACGTGTCATCCTGCC<br>R:GGCAGGATGACACGTGGATGCCAGCACGGATGTGT |
| Fig S3h | **Reference allele primer**<br>F:ATTTCTCTATCGATAGGTACCCAGAGAAGCGCAGCAGCG<br>R:ACTTAGATCGCAGATCTCGAGCGCACAGTGATTCCCGAAGT |

| | |
|---|---|
| | **Primers for mutagenesis**<br>rs78861357 F:GAGACAGCCCTCCCGTTCCGATCTTCTAATC<br>R:GATTAGAAGATCGGAACGGGAGGGCTGTCTC<br>rs76713558 F:GTGTTAGTGAGGGGCTAGACAAGATG<br>R:CATCTTGTCTAGCCCCTCACTAACAC<br>rs79658522 F:GAGCTAGACAAGGTGAGCACGTGAG<br>R:CTCACGTGCTCACCTTGTCTAGCTC<br>rs76525951 F:CACCTGACCCTCATTGCGGCTGCTGG<br>R:CCAGCAGCCGCAATGAGGGTCAGGTG<br>rs79683221 F:GCCTGCATGCCTGGCCGCAGGGCCG<br>R:CGGCCCTGCGGCCAGGCATGCAGGC<br>rs78185801 F:GGGAACGATGCCATCTGCTCGTCTG<br>R:CAGACGAGCAGATGGCATCGTTCCC |
| Fig S3j | **Reference allele primer**<br>F:ATTTCTCTATCGATAGGTACCCCCGGGGGTACTGATGCC<br>R:ACTTAGATCGCAGATCTCGAGAAGATGCTTCCTAGGCTGCGG |
| | **Primers for mutagenesis**<br>rs74785791 F:CAGGTTCTCCTTCCCAGTCCCATGTTC<br>R: GAACATGGGACTGGGAAGGAGAACCTG<br>rs74366004 F:GAACATAAAAACCATACAACTGCAAAGCCTCTGC<br>R:GCAGAGGCTTTGCAGTTGTATGGTTTTTATGTTC<br>rs77702926 F:CACCCTGCCCCACATTTCCCCTGCAC<br>R:GTGCAGGGGAAATGTGGGGCAGGGTG<br>rs78573577 F:GCAAGGATGTCACATGCCTGCAAAGGC<br>R:GCCTTTGCAGGCATGTGACATCCTTGC<br>rs76129108 F:CATGCTTGCTGCCCACTCCGGAGCCCAG<br>R:CTGGGCTCCGGAGTGGGCAGCAAGCATG<br>rs78523927 F:GGCACCACTGTACCGGGTTCTATGAC<br>R:GTCATAGAACCCGGTACAGTGGTGCC |
| Fig S3k | **Reference allele primer**<br>F:ATTTCTCTATCGATAGGTACCAAACGATTGAAACTGGGCCA<br>R:ACTTAGATCGCAGATCTCGAGACTGTGGACGCCCCTCCC |
| | **Primers for mutagenesis**<br>rs11764937 F:GGGGGAGGGTTTTGGGGGGAGATGC<br>R:GCATCTCCCCCCAAAACCCTCCCCC<br>rs79808627 F:GAGCCTCCAGAATCTTTCATCGCCGC<br>R:GCGGCGATGAAAGATTCTGGAGGCTC<br>rs56048221 F:GTTTCATCGCCGCCACAACAAACTCAGG<br>R:CCTGAGTTTGTTGTGGCGGCGATGAAAC |
